# Effective *in vivo* binding energy landscape illustrates kinetic stability of RBPJ-DNA binding

**DOI:** 10.1101/2023.12.19.572376

**Authors:** Duyen Huynh, Philipp Hoffmeister, Tobias Friedrich, Kefan Zhang, Marek Bartkuhn, Francesca Ferrante, Benedetto Daniele Giaimo, Rhett Kovall, Tilman Borggrefe, Franz Oswald, J. Christof M. Gebhardt

## Abstract

Transcription factors (TFs) such as the central DNA-binding hub in Notch signal transduction, RBPJ, bind to specific DNA sequences to regulate gene transcription. How the efficiency of gene regulation depends on the TF-DNA binding kinetics and cofactor interactions is mostly unknown. We determined the DNA binding kinetics and the transcriptional activity of RBPJ and several mutant variants by live-cell single-molecule tracking and reporter assays, and measured their genome-wide chromatin occupation by ChIP-Seq. We observed that cofactor binding, in addition to DNA binding, was required for target site specificity. Importantly, the target site search time of RBPJ was longer than its residence time, indicating kinetic rather than thermodynamic binding stability. Impaired DNA binding, e.g. by mutation K195E related to Adams-Oliver-Syndrome, modulated not only dissociation, but also association to target sites. Impaired cofactor binding mainly altered the rates of unspecific binding and target site association. For other TFs, we also observed longer search than residence times, indicating that kinetic rather than thermodynamic stability of DNA binding might be a general feature of TFs *in vivo*. We propose that an effective *in vivo* binding energy landscape of TF-DNA interactions constitutes an instructive visualization of TF-DNA binding kinetics and the changes upon mutations.

## Introduction

Transcription factors (TFs) such as the DNA-binding hub RBPJ in Notch signal transduction are vital for the regulation of gene expression as they identify specific DNA target sequences and trigger events culminating in gene repression or activation. The efficiency of gene regulation depends among others on the time the TF needs to find its target sequence (target site search time) ^1,2^ and the time it stays bound to the target sequence (residence time) ^1–8^. To which of its potential target sites a TF associates relies on various traits including the accessibility of the site and the presence of other TFs or cofactors ^9–11^, wherefore TFs only occupy a subset of their target sequences ^12,13^. Mutations of the TF affecting its activation or repression capability may give rise to altered DNA binding kinetics. While a mutation within the DNA binding interface is expected to alter the residence time of a TF, it has been observed that such mutations are able to affect the target site search time of a TF ^1,2,14,15^. Therefore, predicting the mechanism of action of a mutated TF from the position of the mutation within the domain-structure of the TF is challenging.

Notch signaling is a short-range cell-to-cell communication pathway in metazoan species evolutionarily conserved from *C. elegans* to *H. sapiens* ^16^. It plays a pivotal role in both embryonic development and adult tissue homeostasis ^17^. Transcription regulation in the Notch signaling pathway is mediated by the DNA binding factor RBPJ (recombination signal binding protein for immunoglobulin kappa J region), which functions as the most important DNA binding hub ^18^. Its DNA binding interface comprises an N-terminal (NTD) and a beta-trefoil (BTD) domain (Supplementary Figure 1). The BTD and the C-terminal domain (CTD) are involved in the interaction with cofactors ^19^. In the absence of Notch signaling, RBPJ functions as a repressor of Notch target genes by recruiting specific corepressor components such as SHARP (SMRT/HDAC1-associated repressor protein) ^20–22^ to form a transcription repressor complex. With active Notch signaling, the Notch Intracellular Domain (NICD1, hereinafter referred to as NICD) is released from the membrane by γ -secretase cleavage and translocates into the nucleus ^23^, where it forms a transcription activation complex with the cofactor MAML ^24,25^. The cofactor composition therefore determines whether the complex around DNA-bound RBPJ acts as repressor or activator of Notch signaling.

A deregulated Notch signaling pathway is responsible for severe congenital diseases such as Alagille Syndrome ^26^ or Adams-Olivier-Syndrome (AOS) ^27^. AOS is associated with the lysine to glutamic acid mutation K195E in the DNA binding interface of RBPJ ^27,28^. Numerous questions related to the signal transfer mechanism of Notch signaling remain. For example, it is unclear whether RBPJ is sufficient to identify specific target sites on chromatin, or whether cofactors are involved in target site selection. Further, it is unknown how cofactors or mutations such as K195E contribute to or alter the DNA binding characteristics of RBPJ *in vivo*.

Here, we determined the target site search time, the DNA residence time and the transcriptional activity of RBPJ and of several DNA and cofactor binding mutant variants by live-cell single-molecule tracking and *in vivo* reporter assays, and measured their genome-wide chromatin occupation by ChIP-Seq. Our measurements revealed important insight into the DNA binding kinetics and specificity of RBPJ and other TFs.

## Results

### Both DNA and cofactor binding contribute to the chromatin binding specificity of RBPJ

To characterize cofactor-dependent DNA binding of RBPJ, we introduced various mutations into the DNA or cofactor binding interfaces. To disturb DNA binding, we chose the mutation R218H (R/H) ^29^, the DNA binding mutation K195E (K/E) found in patients with AOS ^28^, or the triple mutation K195E/R218H/S221D (KRS/EHD) combining the phosphomimetic S221D ^30^ with R/H and K/E ^28^(Figure 1a and Supplementary Figure 1). To disturb cofactor binding, we chose the double mutation F261A/L388A (FL/AA) ^22^, which affects SHARP binding, and the triple mutation.

**Figure 1:**
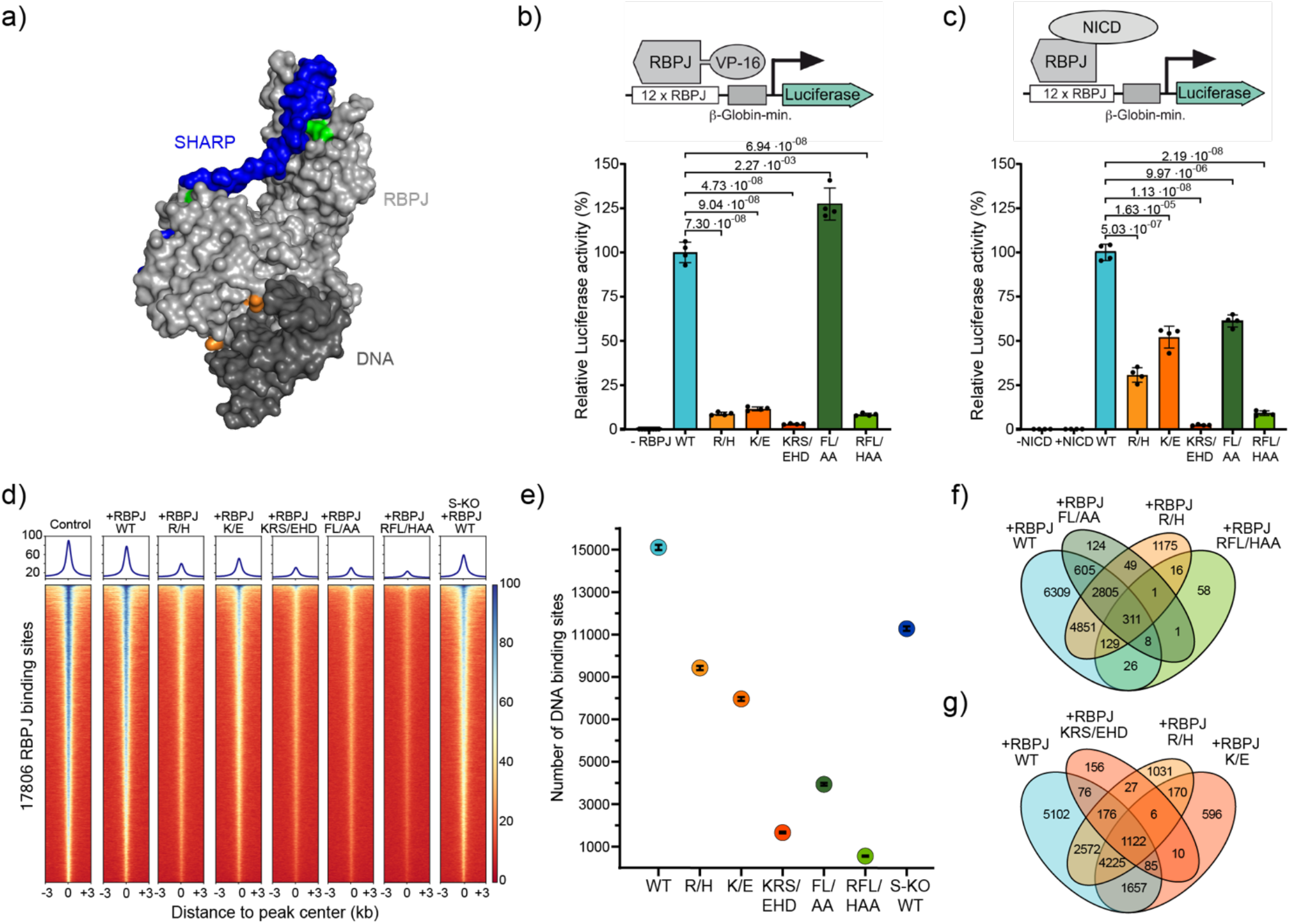
Interactions of RBPJ with cofactors and DNA determine transactivation activity and binding specificity. **a)** Surface representation of a RBPJ-SHARP complex structure (light grey/blue) bound to DNA (dark grey) (PDB entry: 6DKS). Amino acid mutations in interfaces of cofactor binding (green) and DNA-binding (orange) are highlighted. **b), c)** *In vivo* luciferase activity assays. Relative luciferase activity of RBPJ-specific reporter constructs in RBPJ knockout cell line transfected with b) RBPJ-VP16 fusion variants or c) RBPJ variants and co-transfected with NICD (mean values ± s.d.). P-values were determined using two-tailed, unpaired Student’s t-test. Insets: Scheme of the reporter constructs and activating moieties. For details of variants see explanation in the text. - RBPJ: control with untransfected RBPJ knockout cell line, -/+NICD: control with untransfected / NICD-transfected RBPJ knockout cell line. **d)** Heatmap of ChIP-seq reads from RBPJ knockout cell line transduced with HT-RBPJ variants with reads centered around RBPJ binding sites called in an untransduced HeLa cell line control. ChIP performed with an antibody against RBPJ. Read number is color-coded. For details of variants see explanation in the text. S-KO: ChIP-seq reads from SHARP knockout cell line transfected with HT-RBPJ. **e)** Number of binding sites called for HT-RBPJ variants in RBPJ knockout cell line or SHARP knockout cell line. Error bar denotes square root of the value. **f), g)** Venn diagrams depicting common binding cites of indicated HT-RBPJ variants. Source Data are provided as a Source Data file for Figure 1b, c, e.

R218H/F261A/L388A (RFL/HAA), which disrupts both, DNA- and cofactor binding Since SHARP is an important cofactor within the repressor complex of RBPJ, we also assessed how the absence of SHARP influenced RBPJ binding. We first tested the ability of RBPJ variants to functionally bind to the canonical RBPJ target sequence a/GTGGGAAa ^31^. Therefore, we fused the RBPJ variants to the transcription activation domain VP16 ^32^ and utilized a luciferase-based transcription reporter assay in a RBPJ-depleted HeLa cell line, clone #42 ^33^ (Figure 1b and Methods). As expected, the mutations in the DNA binding interface reduced the luciferase signal to either low (R/H, K/E and RFL/HAA) or very low (KRS/EHD) values above the background (Figure 1b). In contrast, solely disturbing the cofactor binding interface (FL/AA) increased the transcription activity, presumably due to impeded binding of corepressor SHARP. This suggests that an intact DNA binding interface is required for functional binding of RBPJ to its canonical site.

To further assess the role of cofactor binding on the transcription activity of RBPJ, we repeated our luciferase assay in the presence of NICD as coactivator, instead of VP16 (Figure 1c and Methods). As expected, mutation FL/AA reduced the transcriptional activity compared to RBPJ-WT. Surprisingly, low transcription activation in the presence of the DNA binding mutations R/H and K/E could be partially rescued by NICD. This was not possible for the triple mutation KRS/EHD or the mutation RFL/HAA that disturbs both DNA and cofactor binding. Thus, the cofactor NICD activates transcription more than VP-16 in this context, if DNA binding is only slightly disturbed.

Next, we tested chromatin binding of RBPJ variants on a genome-wide scale. For later visualization in live cells, we fused wild type RBPJ (RBPJ-WT) and the RBPJ mutants to an N-terminal HaloTag (HT) ^34^. The HaloTag did not interfere with NICD interaction and HT-RBPJ variants showed transcription activity comparable to untagged variants (Supplementary Figure 2). We introduced HT-RBPJ variants into the RBPJ-depleted HeLa cell line clone #42 by lentiviral transduction (Supplementary Figure 3 and Methods), and HT-RBPJ-WT into a SHARP-depleted cell line (clone #30, Supplementary Figure 4). ChIP-Seq reproduced the core binding motif of RBPJ in control cells and confirmed the absence of RBPJ binding in RBPJ-depleted cells (Supplementary Figure 5a-c). We further revealed the binding profile of HT-RBPJ variants in RBPJ-depleted cell lines for RBPJ target genes (Supplementary Figure 5d and confirmed normal RBPJ binding in SHARP-depleted cells (Supplementary Figure 5e-f).

HT-RBPJ-WT closely reproduced the binding profile of endogenous RBPJ (Figure 1d). In accordance with compromised DNA binding, the DNA binding mutations R/H, K/E and KRS/EHD reduced the DNA binding of RBPJ genome-wide (Figure 1d). Moreover, they reduced the number of identified binding sites of HT-RBPJ-WT by ∼38%, ∼47% and ∼89%, respectively (Figure 1e).

Surprisingly, the mutations FL/AA in the cofactor binding interface also reduced the intensity of RBPJ binding (Figure 1d) and reduced the number of binding sites by ∼74% (Figure 1e). Accordingly, for the triple mutation RFL/HAA, almost no binding sites were called, and in the absence of SHARP the number of RBPJ sites identified was also reduced. The position of observed sites in all RBPJ mutants largely overlapped with those of HT-RBPJ-WT, with less than ∼15% *de novo* binding to off-target sites (Figure 1f and g). Taken together, our ChIP-Seq data indicate that not only DNA binding but also cofactor binding of RBPJ plays a key role in specifying chromatin binding of this transcription factor to a majority of its target sites.

### The DNA residence time correlates with transcriptional activity

Next, we quantified the dissociation rate of HT-RBPJ-WT from chromatin by live-cell single-molecule tracking in the RBPJ-depleted HeLa cell line clone #42 (Figure 2b). We visualized individual HT-RBPJ-WT molecules by covalently labeling the HaloTag with HT-SiR dye ^35^, excited fluorescence using HILO illumination to optimize the signal-to-noise ratio ^36^, and detected and tracked fluorescent molecules using the program TrackIt ^37^. We selected cells for imaging that exhibited comparable molecule counts (Supplementary Figure 6).

**Figure 2:**
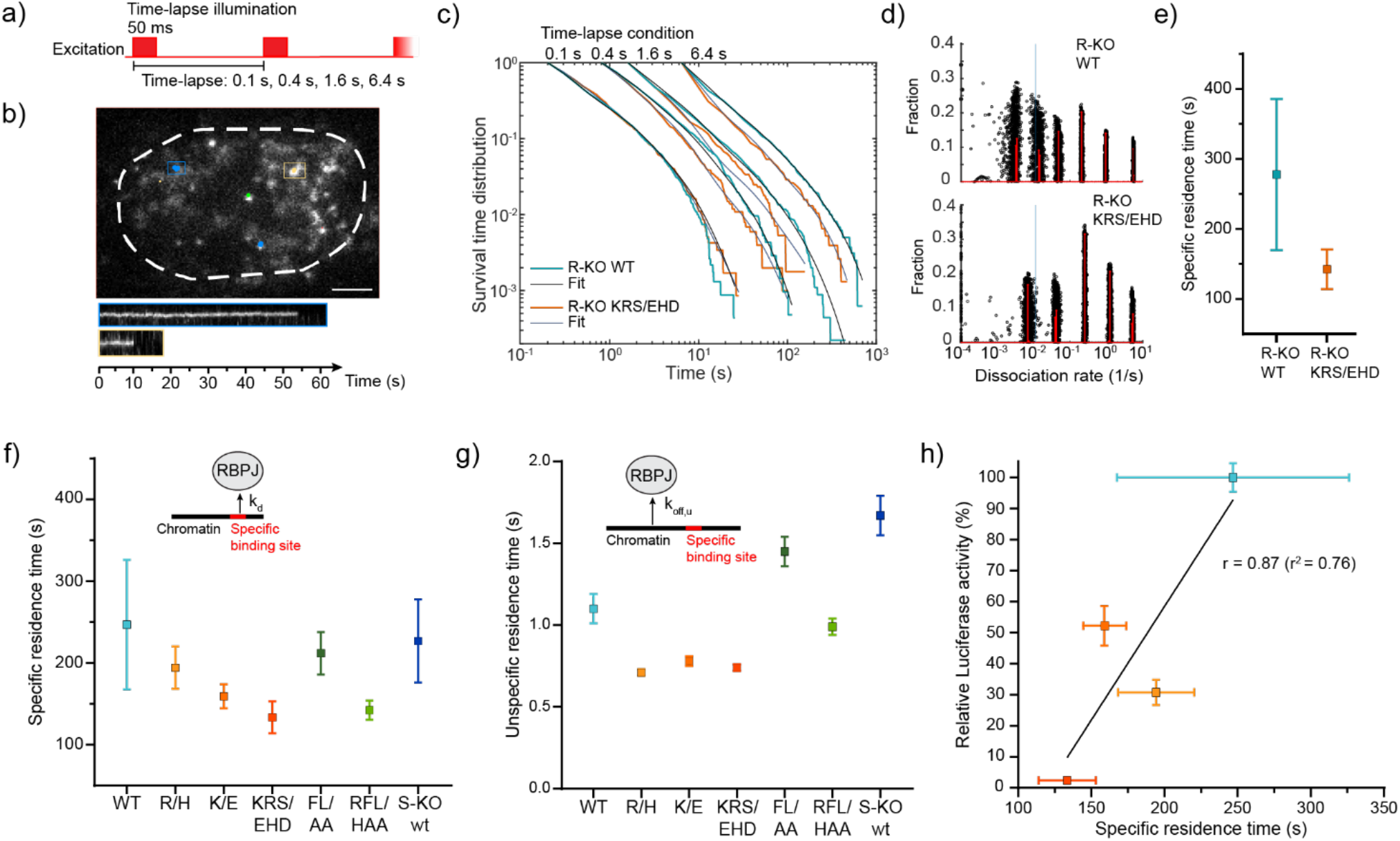
Residence times of HT-RBPJ variants. **a)** Scheme of illumination pattern in time-lapse measurements with indicated camera integration and frame cycle times. **b)** Tracks of single HT-RBPJ-wt molecules overlaid with example image of a 100 ms time-lapse movie (Supplementary Movie 1) and kymographs of indicated molecules. Scale bar is 4 μm. **c)** Survival time distributions of HT-RBPJ-wt (light blue lines) and HT-RBPJ-KRS/EHD mutant (orange lines) at time-lapse conditions shown on top and survival time function obtained with GRID (black lines). For experimental statistics see Supplementary Table 3. **d)** State spectra of dissociation rates of HT-RBPJ-wt and HT-RBPJ-KRS/EHD obtained with GRID using all data (red). As an error estimation GRID was run 500 times with each run using 80 % of the data (black circles). The blue line indicates the dissociation rate of 0.01 s^-1^. For experimental statistics see Supplementary Table 3. **e)** Specific residence times of HT-RBPJ WT and the DNA triple mutant KRS/EHD in Hela RBPJ knock-out cells. **f)** Specific residence times of HT-RBPJ variants (inverse of *k*_*d*_, the weighted average of dissociation rates below 0,01 s^-1^) (Supplementary Table 2). Error bars denote s.d. of the resampled data. Inset: sketch of RBPJ dissociation from a specific binding site with rate constant *k*_*d*_. **g)** Unspecific residence times of HT-RBPJ variants (inverse of *k*_*off,u*_, the weighted average of dissociation rates above 0.01 s^-1^) (Supplementary Table 2). Error bars denote s.d. of the resampled data. Inset: sketch of RBPJ dissociation from an unspecific site with rate constant *k*_*off,u*_. **h)** Relative luciferase activity of HT-RBPJ variants in the presence of NICD versus their specific residence time. Color code as in e). Pearson’s correlation coefficient calculated for HT-RBPJ-wt and DNA binding mutants without considering variants with mutated cofactor binding interface (white squares).Source Data are provided as a Source Data file for Figure 2c-h.

To measure the dissociation rate, we applied several repetitive time-lapse illumination schemes, each comprising an image of 50 ms camera integration time and a time-lapse-specific frame cycle time between 0.1 s and 6.4 s (Figure 2a) ^33^. This approach allows correcting for tracking errors and photobleaching of fluorophores, and ensures covering a broad temporal bandwidth ^37^. We identified binding events as prolonged persistence of fluorescently labeled molecules (Figure 2b, Supplementary Movie 1, and Methods) and collected the fluorescence survival times in histograms (Figure 2c). By performing an inverse Laplace transformation using the GRID method ^38^, we extracted both the event and the state spectrum (Figure 2d) of dissociation rates from these survival time distributions. The event spectrum informs on the number of bound molecules per time showing a certain dissociation rate, while the state spectrum informs on the fraction of bound molecules within a snapshot of time. The dissociation rate spectrum of HT-RBPJ-WT consisted of several distinct rate clusters, corresponding to different residence time classes on chromatin (Figure 2d).

To gain more information on the possible origin of the dissociation rate clusters of HT-RBP-WT, we also measured the dissociation rate spectrum of HT-RBPJ-KRS/EHD (Figure 2c, d). Since this RBPJ variant showed minimal transcriptional activity in both luciferase assays (Figure 1b, c), it should barely bind specifically to DNA. We therefore reasoned that the dissociation rate cluster of HT-RBPJ-KRS/EHD corresponding to the longest residence time could serve to sort dissociation of HT-RBPJ variants into dissociation from either unspecific or specific binding sites (Figure 2d). We then calculated the average specific and unspecific residence times for HT-RBPJ-WT and -KRS/EHD from the inverse of the weighted sum of the corresponding dissociation rates. For HT-RBPJ-WT, we determined unspecific (τ _*u*_) and specific (τ _*s*_) residence times of (0.9 ± 0.1) s and (277 ± 108) s, respectively (Supplementary Table 1-3).

We next characterized to which extent the various mutations in the DNA or cofactor binding interface altered the dissociation of RBPJ from chromatin. To be able to compare the results with those in a SHARP-depleted cell line that still expressed endogenous RBPJ, we stably inserted the HT-RBPJ variants into HeLa cells containing endogenous RBPJ. All tagged variants showed a predominant nuclear localization as expected from endogenous RBPJ and comparable overexpression, which we quantified to be 88% of the expression level of endogenous RBPJ for HT-RBPJ-WT (Supplementary Figure 7a-d). Control measurements revealed comparable residence times of HT-RBPJ-WT or HT-RBPJ-KRS/EHD in presence and absence of endogenous RBPJ (Figure 2e, f).

As expected from compromised DNA binding, the mutations in the DNA binding interface reduced the specific residence times τ_*s*_ on chromatin of (247 ± 79) s for HT-RBPJ-WT to (194 ± 26) s (R/H), (159 ± 15) s (K/E), (133 ± 20) s (KRS/EHD), and (142 ± 12) s (RFL/HAA), respectively (Figure 2f, Supplementary Figure 8 and 9, and Supplementary Table 1-3). The mutations R/H, K/E and KRS/EHD additionally decreased the unspecific residence time of HT-RBPJ-WT (Figure 2g). In contrast, the cofactor binding mutations FL/AA or absence of the cofactor SHARP did not alter the average specific residence time of HT-RBPJ-WT (Figure 2f), but increased the unspecific residence time (Figure 2g). We observed a positive correlation of the specific residence time with the transcriptional activity for the RBPJ variants in which DNA binding was disturbed (Figure 2h), similar to other TFs ^1–8^. This highlights the importance of the DNA residence time for the functioning of TFs.

### The target site search time anticorrelates with transcriptional activity

We next quantified to which extent mutations in the DNA or cofactor binding interface altered the association of RBPJ to a specific target site. In the nucleus, direct association of a TF to a specific target site is slow and association rather proceeds via a faster search mechanism of facilitated diffusion, which combines three-dimensional diffusion in the nucleoplasm and one-dimensional sliding along unspecific DNA ^39^ (Figure 3a). This model entails a transition of the TF from unspecific association to specific binding while sliding over a specific target sequence, potentially associated with a conformational switch ^40,41^. Indeed, structural data from DNA-bound vs unbound Su(H) ^42^, the RBPJ protein in *D. melanogaster*, revealed distinct conformational changes and a reduction of disordered amino acids when bound to its target site (PDB: 5E24, Supplementary Figure 10). Therefore, we assumed that the search process of RBPJ to find a specific target site also follows the mechanism of facilitated diffusion.

**Figure 3:**
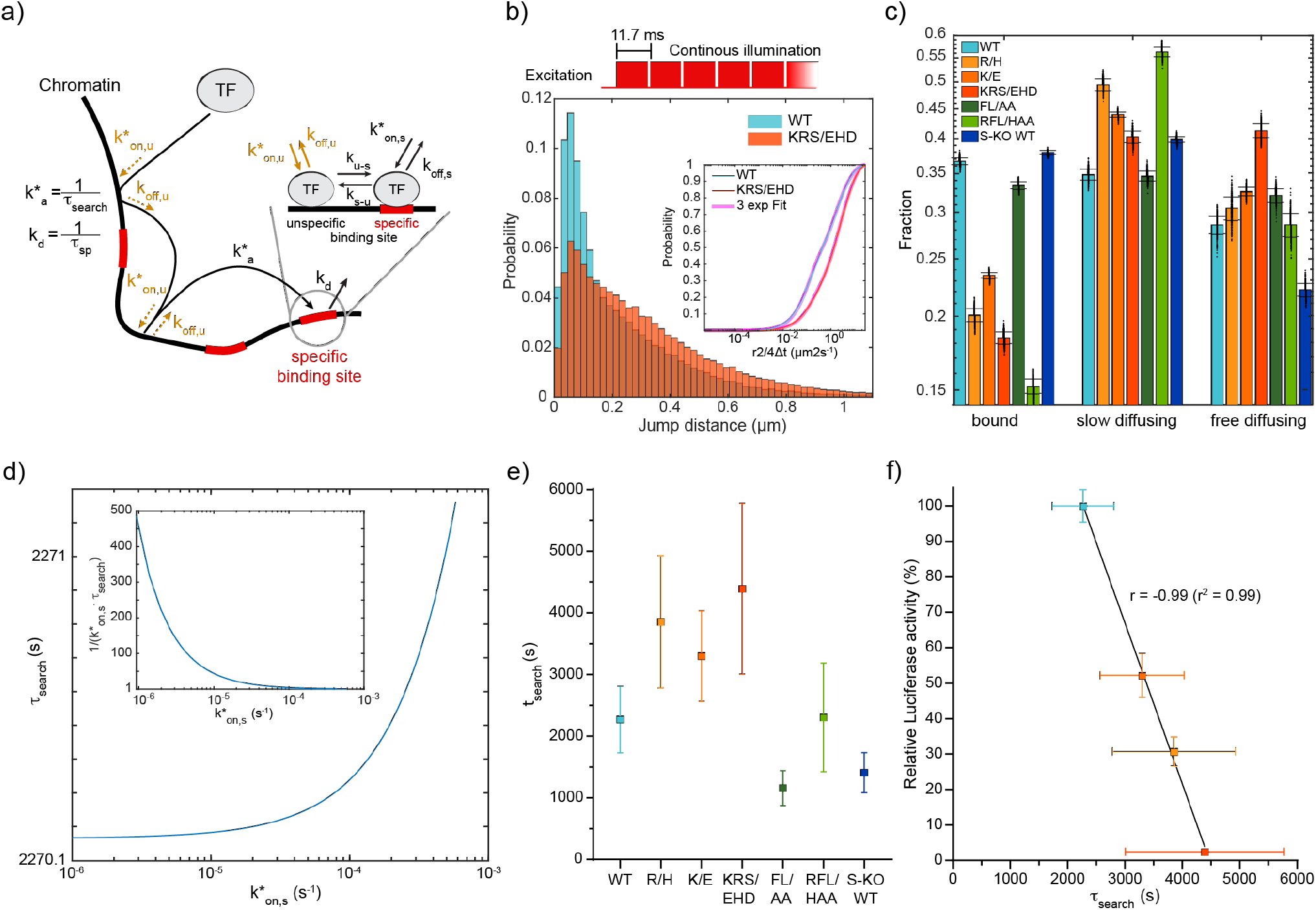
Bound fractions and target site search times of HT-RBPJ variants. **a)** Three-state model of the transcription factor target site search process via facilitated diffusion. Unbound transcription factors associate to a specific binding site with the overall association rate *k\**_*a*_ combining the two pathways of direct association with rate *k\**_*on,s*_ and of indirect association via unspecific binding with rate *k\**_*on,u*_ close to the specific site and sliding into specific binding with the transition rate *k*_*u-s*_. Multiple unspecific binding events may occur before the specific site is bound. Dissociation from the specific binding site occurs with the overall dissociation rate *k*_*d*_ combining the two pathways of direct dissociation with rate *k*_*off,s*_ and indirect dissociation via transition to unspecific binding with rate *k*_*s*-u_ and unspecific dissociation with rate *k*_*off,u*_. **b)** Upper panel: Scheme of illumination pattern for fast tracking with indicated frame cycle time. Lower panel: jump distance distribution of HT-RBPJwt (blue) and HT-RBPJ-KRS/EHD (orange). Inset: cumulative jump distance distributions with three-component diffusion model (pink). **c)** Fractions of the three-component diffusion model and assignment to bound, slow and fast diffusing molecules. Data represents mean values ± s.d. from 400 resamplings with randomly selected 80 % of the data. For experimental statistics see Supplementary Table 5. **d)** Target site search time (*τ*_*search*_) of HT-RBPJ-wt as function of the direct association rate (*k\**_*on,s*_). Inset: fold-acceleration of association via facilitated diffusion over direct association as a function of *k\**_*on,s*_. **e)** Target site search time of HT-RBPJ variants at similar ratio of direct to overall dissociation. **f)** Relative luciferase activity of HT-RBPJ variants in the presence of NICD versus their search time. Color code as in e). Pearson’s correlation coefficient calculated for HT-RBPJ-wt and DNA binding mutants without considering variants with mutated cofactor binding interface (white squares). Source Data are provided as a Source Data file for Figure 3b, c, e, f.

Recently, we devised a three-state model of the search process by facilitated diffusion that relates the target site search time, given by the average time a TF molecule needs to find a specific target sequence among a myriad of unspecific binding sites, to the transition rates between the unbound state and the unspecifically and specifically bound states ^43^ (Figure 3a). The target site search time τ_*search*_ is the inverse of the association rate *k*_*a*_, which combines the pathways of direct association to the specific target site and indirect association via unspecific binding (Methods). It depends on the experimentally accessible parameters of the unspecifically (*p*_*u*_), specifically (*p*_*s*_) and unbound (*p*_*f*_) fractions of the TF, the dissociation rate *k*_*off,u*_ from an unspecific site and the effective specific dissociation rate *k*_*d*_. *k*_*d*_ combines the pathways of direct dissociation from the specific target site and indirect dissociation via unspecific binding. In addition, one of the direct target site association (*k*_*on,s*_) or dissociation (*k*_*off,s*_) rates or of the microscopic transition rates *k*_*u-s*_ or *k*_*s-u*_ between unspecific and specific binding needs to be known to calculate the other missing parameters and the target site search time ^43^ (Methods).

To obtain the fraction of molecules bound both unspecifically and specifically, we measured the diffusive behavior of HT-RBPJ variants by recording continuous movies with 10 ms camera integration time (Figure 3b, Supplementary Table 4 and 5, and Supplementary Movie 2). We collected the jump distances of detected single-molecule tracks in cumulative histograms and analyzed these with a diffusion model including slow, intermediate and fast diffusing molecules, which described data better than a two-component model (Figure 3b, Supplementary Figure 11 and Methods) ^37,44^. The diffusion coefficients of HT-RBPJ variants were similar, compatible with their similar size (Supplementary Figure 12a). The slow diffusion component arises due to slow movement of chromatin bound by HT-RBPJ molecules and the single-molecule localization error. It represents the fraction of molecules bound to either unspecific or specific chromatin sites ^45–47^. We found a bound fraction of ∼37% for HT-RBPJ-WT, similar to the value of 34% observed in *D. melanogaster* salivary gland cells ^48^. A linear relation between bound molecules and all detected molecules confirmed that binding was not saturated in our overexpression conditions (Supplementary Figure 7f). As expected, the overall bound fraction was reduced to ∼20% (R/H), ∼23% (K/E) and ∼18% (KRS/EHD) if the DNA binding interface of RBPJ was disturbed (Figure 3c and Supplementary Table 4 and 5). Moreover, the fraction of specifically bound HT-RBPJ molecules correlated with the number of target sites identified by ChIP-Seq, similar to previous findings (Supplementary Figure 12b) ^49^.

We calculated the target site search time τ _*search*_ of HT-RBPJ-WT by combining the results of our residence time and binding fraction measurements and varying the kinetic rates *k\**_*on,s*_, *k*_*off,s*_, *k*_*u-s*_ and *k*_*s-u*_ that were not experimentally accessible (Methods). This yielded a minimal target site search time of ∼2270 s, which increased in a small interval up to ∼2271 s for a variation of *k\**_*on,s*_ over three orders of magnitude (Figure 3d, Supplementary Figure 13, and Supplementary Table 6). At the upper limit of τ_*search*_, the time of direct association equaled the target site search time, and a description of the search mechanism by facilitated diffusion starts to break down.

To compare the target site search times of different HT-RBPJ variants, we kept the ratio of the direct dissociation rate *k*_*off,s*_ to the measured effective dissociation rate *k*_*d*_ constant for all HT-RBPJ variants All mutations in the DNA binding interface increased the target site search time of HT-RBPJ-WT (Figure 3e and Supplementary Table 7). In contrast, the mutations FL/AA disturbing cofactor binding or the absence of SHARP did not alter the search time. Interestingly, the target site search time of variants with disturbed DNA binding domain anticorrelated with transcriptional activity (Figure 3f). Thus, in addition to the DNA residence time, the target site search time is important for RBPJ function.

### RBPJ function is governed by association and dissociation rates

In a binding reaction at equilibrium, the effective association and dissociation rates of the reactants are coupled through the equilibrium constant and define the binding energy of the reaction (Figure 4a and Methods). If the effective association rate is larger than a given dissociation rate, the energy of the bound state is lower than the energy of the unbound state, and the reaction is thermodynamically stable (Figure 4a). If the effective association rate is smaller than the given dissociation rate, the energy of the bound state is higher than the energy of the unbound state. Importantly, in this case, the interactions underlying the adhesion of both reactants are still present, and the reactants are kinetically trapped in the bound state, making it kinetically stable.

**Figure 4:**
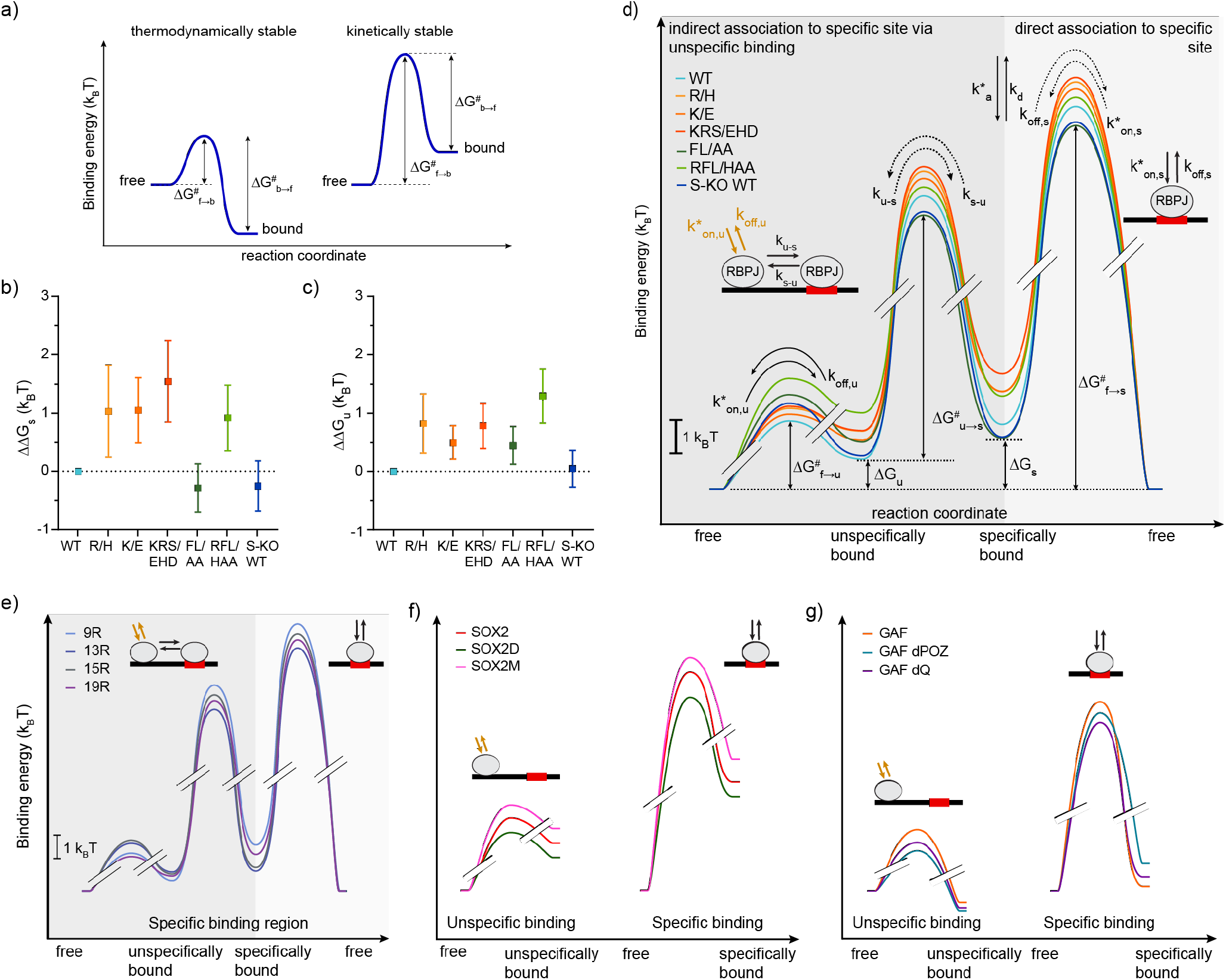
*In vivo* binding energy landscape of transcription factors. **a)** Sketch of binding energy landscapes of a thermodynamically stable and a kinetically stable binding interaction. In both cases, the energy barrier of unbinding and thus the residence time in the bound state is equal. **b)-c)** Binding energy differences. Differences in b) specific binding energies *ΔΔG*_*s*_ and c) unspecific binding energies *ΔΔG*_*s*_ between HT-RBPJ-wt and mutants. **d)** *In vivo* chromatin binding energy landscapes of HT-RBPJ variants. Energy differences between bound states or transition barriers and the free state as well as kinetic rates of the target site search process by facilitated diffusion are indicated. Dark grey shade: specific association via unspecific binding and subsequent transition to specific binding; bright grey shade: direct specific association. Only relative, but not absolute energy differences of transition barriers can be obtained. Dotted arrows indicate undetermined rates, limits of the relative transition barriers are given in Supplementary Figure 9. **e)**-**g)** *In vivo* chromatin binding energy landscapes of e) TALE variants^2^, f) SOX2 variants constructed from published kinetic rates^14^, and g) GAF variants constructed from published kinetic rates^15^. Source Data are provided as a Source Data file for Figure 4a-h.

To illustrate the effect of mutations in the DNA and cofactor binding interfaces of RBPJ on the RBPJ binding kinetics in the nucleus, we propose to construct the effective *in vivo* binding energy landscape of RBPJ. It can be obtained from its unspecific and specific bound fractions, the effective association rate, and the dissociation rate (Methods). Importantly, while TF-DNA binding free energies are typically compared at standard concentrations of binding sites, taking the cellular concentrations of binding sites of the TFs into account enables comparing the situation the TFs actually encounter in the nucleus.

We first determined the effective *in vivo* binding energy Δ G_s_ of specific binding, that is the effective energy difference between binding to a specific target sequence and the unbound state, from the ratio of the specifically bound to the unbound fraction of molecules for each of the HT-RBPJ variants (Supplementary Table 8 and Methods). The contribution of a mutation to the specific effective binding energy of RBPJ is then contained in the energy difference ΔΔG_s_ with HT-RBPJ-WT (Figure 4b). Consistent with compromised DNA binding, all mutations in the DNA binding interface, R/H, K/E, KRS/EHD, and RFL/HAA increased the specific effective binding energy of HT-RBPJ-WT. In contrast, mutation FL/AA disturbing cofactor binding or absence of SHARP slightly decreased ΔG_s_ compared to HT-RBPJ-WT. Thus, the specific effective binding energies of HT-RBPJ variants illustrate mutation-specific changes of the RBPJ chromatin binding stability in the nucleus.

Similar to ΔG_s_ of specific binding, we determined the unspecific effective binding energy ΔG_u_ from the ratio of the unspecifically bound fraction to the unbound fraction of HT-RBPJ variants (Figure 4c) (Methods). Again, all mutations in the DNA binding interface increased the unspecific effective binding energy of HT-RBPJ-WT.

We further calculated other basic points of the effective *in vivo* binding energy landscape of HT-RBPJ-WT and mutants: in addition to ΔG_s_ and ΔG_u_ also the effective energy barriers ΔG^#^_f→u_ and ΔG^#^_f→s_ of the direct unspecific and specific association and dissociation processes, and the effective energy barrier ΔG^#^_u→s_ of the transition from unspecific to specific binding at a target sequence (Figure 4d and Supplementary Table 8). We again used the ratios of bound to unbound fractions as well as the kinetic rates obtained from the three-state model of the target site search process to calculate these effective energy barriers (Methods). While we could not obtain the absolute height of the barriers without knowledge of the frequency factor, the relative height of barriers can be compared. The binding energy landscape of HT-RBPJ-WT was comparable in the presence and absence of endogenous RBPJ (Supplementary Figure 14).

A striking feature of the effective *in vivo* binding energy landscapes of HT-RBPJ variants is that the specifically bound state has a higher energy than the unspecifically bound and the unbound, free state, ΔG_s_ > ΔG_u_ > 0 (Figure 4d). Counterintuitively, this means that binding of RBPJ to a specific target sequence is thermodynamically unstable *in vivo*, whereas unspecific binding and even more the free state are thermodynamically favored. This arises from the longer association time than residence time of unspecific and specific binding, and is reflected in the small bound fractions below 0.4 of HT-RBPJ variants (Figure 4a and 3c). Yet we emphasize that binding of RBPJ to chromatin is kinetically stable, with residence times of up to ∼250 s for HT-RBPJ-WT.

### Transcription factor binding is kinetically, not thermodynamically stable *in vivo*

Encouraged by the insight obtained from the effective *in vivo* binding energy landscape of RBPJ, we reanalyzed our previously measured live-cell single molecule data of TALE (transcription activator like effector) TFs (TALE-TFs) ^2^. The TALE-TFs differed in the length of their DNA binding domain and showed varying residence times on chromatin ^2^. We now additionally calculated the basic points of the effective binding energy landscape of TALE-TFs from the bound fractions and kinetic rates analogous to RBPJ (Figure 4e and Supplementary Table 9). Similar to RBPJ, binding of TALE-TFs to chromatin is not thermodynamically but kinetically stable *in vivo*. Moreover, we calculated the effective binding energy landscapes of SOX2 and GAF and of various DNA binding domain- or activation domain-mutants of these factors using published kinetic rates ^14,15^ (Figure 4f and g, Supplementary Table 10). Analogous to RBPJ and TALE-TFs, specific chromatin binding of SOX2 and GAF occurred at higher effective energy than unspecific binding and the unbound state and thus was thermodynamically unstable *in vivo*. This indicates that kinetic instead of thermodynamic stability of chromatin binding might be a general feature of transcription factors *in vivo*.

## Discussion

AOS is linked to the autosomal dominant mutation K195E (K/E) in the DNA binding interface of RBPJ ^27,28^. Our live-cell measurements showed that HT-RBPJ-K/E still binds to DNA and is able to activate gene transcription in presence of NICD, albeit to a lower degree than HT-RBPJ-WT (Figure 1c). In particular, we found that mutation K/E increased the target site search time of RBPJ 1.5-fold, reduced the residence time of RBPJ at specific target sites by ∼36% and increased the effective binding energy by ∼1 k_B_T, corresponding to a 2.1-fold smaller specifically bound fraction. ChIP-Seq experiments confirmed reduced chromatin binding of HT-RBPJ-K/E, and revealed a loss of 47% of the HT-RBPJ-WT target sites (Figure 1e). Our findings allow refining possible molecular mechanisms underlying AOS: If one allele carries the mutation, the amount of wild-type RBPJ will be reduced, thus reducing the occupation frequency of target sites exclusively bound by RBPJ-WT. This reduced target site occupation and the shorter residence time and lower transcription activation potency of RBPJ-K/E at sites bound by both species should limit the repression and activation of genes, in addition to proposed cofactor sequestration ^28^. Moreover, off-target binding of HT-RBPJ-K/E as revealed by ChIP-Seq (Figure 1g) might lead to a gain-of-function phenotype. This scenario could be similar as observed for point mutations of IRF4 associated with autosomal dominant combined immunodeficiency ^50,51^. Overall, our data predict de-repression and lower than normal Notch-activated transcription of numerous Notch target genes in AOS.

The effective *in vivo* binding energy landscape of RBPJ reveals important insight about the mechanism of action of RBPJ mutations (Figure 4d). The DNA binding mutations R/H, K/E, and KRS/EHD weakened both specific and unspecific binding. Binding was destabilized not only by increasing the direct dissociation rates *k*_*off,s*_ and *k*_*off,u*_, but also by decreasing the direct association rates *k\**_*on,s*_ and *k\**_*on,u*_. This resulted in an increase of both the transition barriers ΔG^#^_f→s_ and ΔG^#^_f→u_ and the energies ΔG_s_ and ΔG_u_ of specific and unspecific binding. Interestingly, changes in the barrier ΔG^#^_f→s_ of specific binding were mirrored in the changes of the barrier ΔG^#^_u→s_ to switch from unspecific to specific binding.

In contrast to mutations in the DNA binding interface, the mutations FL/AA in the cofactor binding interface or absence of SHARP mainly increased the barrier ΔG^#^_f→u_ to unspecific binding, while the barriers to specific binding where decreased, resulting in a lower effective binding energy ΔG_s_ (Figure 4d). The mutations FL/AA impede interaction of RBPJ with cofactors including SHARP, RITA1 and NICD ^22,25,52^. As expected, the NICD-mediated transcription activation potency of HT-RBPJ-FL/AA was reduced (Figure 1c). We further observed that HT-RBPJ-FL/AA loses the ability to associate to 75% of the HT-RBPJ-WT target sites, despite an intact

DNA binding domain (Figure 1e). Thus, similar to the role of the activation domain of several TFs in determining DNA specificity ^53,54^, some cofactors of RBPJ might play a role in guiding RBPJ to its cognate target sites.

Our approach to determine the effective *in vivo* binding energy landscape by measuring bound fractions and kinetic rates using single-molecule tracking is model-free. It is therefore not limited to protein-DNA interactions. The model that we additionally employed here for the search process of TFs enabled revealing more details of TF-DNA interactions such as the transition from unspecific to specific binding at the target sequence, but is generally not required. Similar to *in vivo* measurements of protein-protein dissociation constants by single-molecule tracking ^55^, fluorescence cross-correlation spectroscopy ^56–58^ or Förster resonance energy transfer ^59^, our single-molecule approach including kinetic rates should be readily applicable to other biomolecular interactions *in vivo*.

To conclude, our measurements uncovered new insight into the cofactor-dependance of the DNA binding specificity of RBPJ and the molecular mechanism of the disease-related mutation K195E. Moreover, reconstructing the effective binding energy landscapes of various RBPJ variants and other TFs from their bound fractions and association and dissociation rates revealed that TF-DNA binding is thermodynamically unstable *in vivo*, while the slow association and dissociation rates of the TFs ensure kinetic stability. It will be an important task for the future to develop models of gene regulation including slow target site search kinetics of TFs in non-equilibrium conditions ^60^. Moreover, kinetic rather than thermodynamic stability might also be valid for protein-protein and other biomolecular interactions *in vivo*, if association is slow compared to dissociation.

## Material and Methods

### Cell Culture

For all experiments, we used HeLa-derived cell lines, which were grown in DMEM additional supplemented with 10 % fetal bovine serum, 1 % glutamax, 1 % non-essential amino acids and 1% sodium pyruvate and cultured at 37 °C and 5% CO_2_.

### Transient DNA transfection

To transiently transfect HeLa^RBPJ KO^ cells (clone #42) with expression plasmids and reporter plasmids (expression plasmids are listed in Supplementary Table 11) for subsequent luciferase assays, we used the Lipofectamine 2000 transfection reagent (Invitrogen, Cat.No.: 11668019) according to the manufacturer’s instructions.

### Luciferase assay

To determine the luciferase activity of previously transfected HeLa^RBPJ KO^ cells, we used the luciferase assay system from Promega (Promega, Cat. No.: E1501). To do so, we seeded 2.25 × 10^4^ cells per well onto a 48-well plate and transfected them after 24 h with 0.25 μg reporter plasmid per well. Furthermore, were co-transfected some cells with 100 ng of expression plasmids for RBPJ mutants. For cell lysis, we applied 100 μl of 1:5 diluted cell culture lysis reagent (Promega, Cat. No.: E1531) to each well, 24 h after transfection. We centrifuged the resulting cell lysates and applied 10 μl of the supernatant to a 96-well plate. Finally, we used a Luminometer Microplate reader LB960 (Berthold Technologies, Cat. No: S11902) in order to determine the luciferase activity. We performed at least four independent experiments with two technical replicates.

### Co-Immunoprecipitation

To verify protein-protein interaction between Halo-tagged RPBJ proteins and Flag-tagged NICD, we performed Co-Immunoprecipitation experiments. For Co-immunoprecipitation of Halo-tagged RBPJ proteins together with Flag-tagged NICD we co-transfected HEK293 cells with expression plasmids for the given proteins (used plasmids are listed in Supplementary Table 12). Cells were lysed 24 h after transfection using 600 μl CHAPS lysis buffer. We incubated 500 μl CHAPS lysate with agarose beads conjugated with anti-Flag antibodies (Merck, Cat. No.: A2220) over night at 4 °C on an overhead shaker. Samples were washed 6 times with CHAPS lysis buffer and resuspended in 1x Laemmli buffer for subsequent SDS-PAGE.

### Generation of CRISPR/Cas9 depleted cells

We designed the CRISPR/Cas9 guides using the online tool available at http://crispor.tefor.net/. We added the desired 5’ overhangs and phosphorylated, annealed and ligated the oligos into the px459 v2.0 (Addgene #62988)^61^ predigested with BbsI. The *SHARP* depletion was generated with the combination of hSHARP guide #1 and hSHARP guide #2. The *RBPJ* depletion was generated with the combination of hRBPJ guide #1 and hRBPJ guide #2 (sequences of the guides in Supplementary Table 13). We transfected HeLa cells (ATCC CCL-2) with 10 μg of each px459 v2.0 plasmid together with 40 μl of linear PEI (Polyscience 23966) using standard protocols. We selected the cells with puromycin before establishing single cell clones.

### RNA extraction, reverse transcription, quantitative PCR (qPCR)

To purify the total RNA, we used the RNeasy Mini Kit (Qiagen #74104), the QIAshredder (Qiagen #79654) and the DNase I (Qiagen #79254) accordingly to the manufacturer′s instructions. For generation of cDNA, we used 1 μg of RNA and retro-transcribed it using random primers and SuperScript II reverse transcriptase (Invitrogen #18064-014). We assembled qPCR reactions with QuantiFast SYBR Green RT-PCR Kit (Qiagen #204156), gene-specific oligonucleotides (Supplementary Table 14) and analyzed using the LightCycler480™ system (Roche Diagnostics). We calculated mRNA expression levels relative to the housekeeping gene *Hypoxanthine Phosphoribosyltransferase 1* (*HPRT*).

### ChIP-Seq

To prepare HeLa cells for ChIP-seq experiments, we washed the cells twice with PBS, fixed them for 1 hour at room temperature in 10 mM dimethyladipimate (DMA, Thermo Scientific 20660) dissolved in PBS and, after washing once in PBS, crosslinked in 1% FMA for 30 min at room temperature. We blocked the FMA reaction by adding 1/8 volume of 1 M glycine pH 7.5 and incubated for 5 min at room temperature. We performed chromatin immunoprecipitation (ChIP) essentially as previously described ^62^ using an anti-RBPJ antibody (Cell Signaling Technology, 5313S). For spike-in purpowes, we used chromatin from *D. melanogaster* Schneider cells in presence of 2 μg of anti-His2Av antibody (Active Motif 61686) for each immunoprecipitation. Antibodies are listed in Supplementary Table 15.

We prepared libraries using the Diagenode MicroPlex Library Preparation kit v3 (Diagenode C05010001) following the manufacturer’s instructions with few modifications. Subsequently, we purified libraries with Agencourt AMPure XP Beads (Beckman Coulter, #A63881), and quantified and analyzed on an Agilent Tapestation device. Finally, we performed sequencing on a NovaSeq device at Novogene UK.

### ChIP-Seq analysis

We applied quality and adapter trimming to raw sequencing reads with TrimGalore (https://github.com/FelixKrueger/TrimGalore).

Next, we aligned the trimmed reads to the human reference genome (hg19, downloaded from Illumina’s iGenomes) using Hisat2 ^63^ with “-k 1 – no-spliced-alignment –phred33” parameters and stored them as binary alignment maps (BAM). To filter BAM files for PCR duplicates, we used the *MarkDuplicates* function of the Picard tools (available at http://broadinstitute.github.io/picard/) with “REMOVE_SEQUENCING_DUPLICATS = true REMOVE_DUPLICATES = true” parameters. Next, we generated normalized coverage tracks (bigWigs) using the Deeptool’s *bamCoverage* function ^64^ based on the filtered BAM files. To call peaks for the individual samples, we used PeakRanger ^65^ with p- and q-values cutoffs of 0.0001 and the matching input for each sample. We selected only peaks that were detected in both replicates and not overlapping with ENCODE’s blacklisted regions (https://github.com/Boyle-Lab/Blacklist/) for further analysis. By using IGV ^66^, we visually inspected the conclusiveness of the peak calling. Subsequently, we performed motif analysis using the MEME-suite ^67^ based on the summits of RBPJ peaks detected by PeakRanger +/- 50 base pairs. We generated heat maps using Deeptools’s *computeMatrix* and *plotHeatmap* functions based on the RBPJ peak set detected in HeLa control cells and the normalized coverage tracks for all samples. To generate Snapshots for example regions, we used Gviz ^68^.

### Stable cell line generation

We thawed LentiX-293T packaging cells at least one week prior to the virus production. To start the virus production, we transfected LentiX cells with 3 different plasmids, the plasmid pLV-tetO-Oct4 (Addgene #19766)^69^ including the sequence of interest, the packaging plasmid psPAX2 (Addgene #12260) and the virus envelope plasmid pMD2.G (Addgene #12259) using JetPRIME (Polyplus-transfection, #114-15). To obtain a sufficient amount of virus, we let LentiX cells grow to 80 % confluency on a 10 cm culture dish before transfection. Next, we mixed 10 μg of plasmid encoding for the Halo-tagged construct of interest, 7.5 μg psPAX2 and 2.5 μg of pMD2.G with 500 μl of jetPRIME buffer and briefly vortexed the resulting mix. We added 30 μl of jetPRIME transfection reagent to the mixture and shortly vortexed. Subsequently, we incubated the transfection mixture for 10 minutes at room temperature. After incubation, we added the mixture dropwise to the LentiX cells and incubated them for 2 days at 37 °C and 5% CO_2_.

One day before lentiviral transfection, we seeded 40,000 Hela cells in the wells of a 6-Well plate. We harvested the viruses by collecting the supernatant (9-10 ml) of LentiX cells with a syringe, filtering it through a Whatman™ 0.45 μm membrane filter and collecting the flow-through in a 15 ml falcon tube. After that, we added 1 ml of filtered virus solution to the Hela cells followed by an incubation for 3 days at 37 °C and 5% CO_2_.

After 3 days, we expanded the cells on a 10 cm culture dish by washing them once with 1 ml PBS and trypsinizing them with 0.5 ml 0.05% Trypsin for 4 minutes at 37°C and 5% CO_2_. We stopped trypsinization by adding 4.5 ml DMEM.

To test whether the viral transduction was successful, we labeled the cells with 2.5 μM HaloTag-Ligand (HTL) TMR (Promega #G8251) for 15 min, following the manufacturer’s protocol, and observed them under a confocal spinning disk microscope. In order to obtain a highly homogenous positive cell population, we sorted the cells via their fluorescence using FACS. For this, we labeled cells with 1.25 μM HTL-TMR following the manufacturer’s protocol. To adjust the flow cytometer gates, we used unlabeled Hela cells.

### Western Blotting

To verify the stable expression of target proteins in the aforementioned stable cell lines, we performed western blots in order. Therefore, we lysed cells by applying 120 μl of CHAPS lysis buffer containing protease inhibitors to cell pellets. After an incubation on ice for 1 h, we centrifuged the samples for 30 min at 14,000 rpm and 4 °C. Subsequently, we collected the supernatant in a new 1.5 ml reaction tube and determined the protein concentration of each sample by a Bradford protein assay (Biorad, Cat. No.: 5000006). For each sample, we mixed 20 μg of protein with 6x Laemmli buffer and applied it to a 10% SDS polyacrylamide gel. After gel electrophoresis, we blotted the proteins at RT on a PVDF membrane (Merck, Cat. No.: IPVH00010). We blocked the membranes for 1 h at RT with sterile filtered 5 % bovine serum albumin (BSA) (Serva, Cat. No.:9048-46-8) dissolved in TBS with 0.1% Tween-20, prior to incubation with the primary anti-Halo-Tag antibody (Mouse monoclonal antibody, Promega, Cat. No.: G9211). For incubation with the primary anti-RBPJ antibody (Rat monoclonal antibody, Cosmo Bio, Cat. No.: SIM-2ZRBP2), ee blocked the membranes under the same condition as above using 5% skim milk (PanReac AppliChem, Cat. No.: A0830) dissolved in PBS with 0.1% Tween-20. To stain with the anti-Halo-Tag antibody, diluted 1:500 in TBS-T, we incubated the membranes over night at 4 °C. To apply the anti-RBPJ antibody, we used PBS-T for a final dilution of 1:1,000. After washing the membranes three times with PBS-T to remove unbound primary antibodies, we used horseradish peroxidase conjugated secondary antibodies against mouse (GE Healthcare, Cat. No.: NA931V) or rat (Jackson ImmunoResearch, Cat. No.: 112-035-071) to detect primary antibodies. To detect the target proteins, we incubated the membranes for 1 min in 5 ml of ECL solution (Cytiva,Cat. No.: RPN2209). We detected the resulting chemiluminescence signal with High performance chemiluminescence films (Cytiva, Cat. No.: 28906837). All antibodies used are shown in Supplementary Table 16.

### Quantification of Western Blot data

We performed Western blots using five individual lysates of wild type HeLa cells stably expressing HT-RBPJ-WT to quantify the expression of HT-RBPJ-WT proteins relative to endogenous RBPJ. We determined expression levels by quantifying the mean intensities of the bands of endogenous RBPJ and HT-RBPJ-WT using ImageJ (https://imagej.net/ij/). The expression levels of HT-RBPJ-WT are calculated relative to endogenous RBPJ. Finally, we calculated the mean of all individual relative values to determine the final relative expression level of HT-RBPJ-WT.

### Flow cytometry to determine the cellular abundance of HT-RBPJ-WT

To determine the average amount of HT-RBPJ-WT proteins in the HeLa cell line stably expressing HT-RBPJ-WT, we performed flow cytometry (FCM) measurements with an Attune NxT flow cytometer and compared the intensity level with a calibrated reference U2OS cell line stably expressing Halo-tagged CTCF (C32) ^70^. We calibrated the device and determined the background fluorescence using untreated HeLa and U2OS cells. We seeded the cells in 10 cm dishes and stained them on the day of the FCM measurement at a confluence of 80 %. To stain the cells, we incubated them for 30 min at 37 °C with 500 nM Halo-TMR ligand. Subsequently, we aspirated the medium and washed the cells once with PBS. After removing PBS, we added fresh pre-warmed medium before detaching the cells by trypsinization. We transferred the cell suspension into fresh DMEM and determined the cell concentration using a Neubauer hemocytometer. Next, we centrifuged the samples for 5 min at 1,200 rpm at 4 °C. Prior to subjecting the cells to FCM, we aspirated the supernatant and resuspended the cells stored on ice in 220 μl PBS.

To determine the average amount of expressed HT-RBPJ-WT in HeLa cells, we calculated the relative fluorescence intensity by subtracting the background fluorescence and compared it to the reference fluorescence intensity of U2OS C32 cells (Supplementary Figure 15). We estimated a mean protein abundance of 90,863 HT-RBPJ-WT molecules per cell. Since HT-RBPJ-WT was overexpressed 0.88-fold compared to endogenous RBPJ (Supplementary Figure 7d), this corresponded to an average of 102,855 endogenous RBPJ molecules and a total number of 193,718 HT-tagged or endogenous RBPJ molecules per cell. Assuming an ellipsoidal nucleus with volume π/6*8*8*5 μm^3^, the corresponding average RBPJ concentration was ∼1.92 μM.

### Widefield fluorescence microscopy

We used fluorescence microscopy to investigate the subcellular localization of the stably expressed HT-RBPJ variants. Therefore, we seeded 4 × 10^4^ cells of each investigated stable cell line into one well of a 2-well chamber glass cover slip (Nunc LabTek, Cat. No.: 155380) that was previously incubated with a 1x fibronectin solution (Sigma-Aldrich, Cat. No.: F2006) for 30 min at 37 °C. Afterwards, we aspirated the fibronectin solution and subsequently washed the chamber coverslips twice with PBS. After 24 h, we stained the cells with 1 ml of a 1:2,000 TMR Halo-ligand solution (Promega, Cat. No.: G8251) to fluorescently label the Halo-tagged proteins. After an incubation time of 15 min at 37 °C, we removed the Halo-ligand solution and washed the chamber coverslips three times with fresh DMEM medium, followed by an additional application of 1 ml of DMEM and a subsequent incubation of 30 min at 37 °C. We fixed the cells and labeled the DNA by incubating the cells with 1 ml of a DAPI solution (1:10,000 in PBS) on a shaker at RT. Finally, we washed the samples 5 times for 5 min with PBS and added one drop of fluoromount-G mounting medium (SouthernBiotech, Cat. No.: 0100-01) to each well prior to applying a cover slip.

Alternatively, in order to confirm the SHARP knock-out in HeLa SHARP knock-out clones #36 and #30, we used immunofluorescence. We applied 1 ml of DMEM, containing 45,000 cells, on cover slips and incubated them overnight at 37 °C. On the next day, we aspirated the medium and washed the coverslips once with PBS. After fixing the cells, we permeabilized them by applying 1 ml of 0.2 % Triton for 2 min. After two washing steps with PBS, we blocked unspecific binding of antibodies using PBS that contained 1% BSA, 1 % FBS and 0.1 % fish-skin gelatine for 30 min at RT. We diluted the primary antibody (anti-SHARP.1 ^20^) 1:500 in blocking buffer and incubated the cells for 3 h. After five washing steps with PBS, we applied the secondary antibody for 1 h as a 1:1,000 dilution in blocking buffer. After 30 min, we applied DAPI for a final dilution of 1:20,000, which was later removed with the secondary antibody. After 6 final washing steps with PBS, we added a drop of fluoromount-G mounting medium (SouthernBiotech, Cat. No.: 0100-01) on an object slide and placed the coverslip on top of it. Specifications of the used antibodies are shown in Supplementary table 17. To image the cells, we used an Olympus IX71 fluorescence microscope, a 100 W mercury lamp (Osram, HBO 103W/2) and a digital camera (Hamamatsu, C4742-95). To detect TMR (Excitation: ET545/25, Emission: ET 605/70), we used a Cy3 ET filter set (AHF, Cat. No.: F46-004), a suitable filter set for DAPI detection (Excitation: D360/50, Emission: D460/50 and an EGFP ET filter set (Excitation: ET470/40, Emission: ET 525/50) (AHF, Cat. No.: F46-002) for detection of green fluorescence.

### Preparation of cells for single-molecule imaging

One day prior to single-molecule imaging, we seeded cells on a 35 mm heatable glass bottom dish (Delta T, Bioptechs). For time-lapse imaging, we stained the cells with 3-6 pM HTL-SiR on the day of imaging. For that, we incubated cells with HTL-SiR for 15 min at 37 °C and 5% CO_2_ and washed them once with PBS followed by a recovery step of 45 min at 37 °C and 5% CO_2_ in DMEM. Directly before imaging, we washed cells three times with PBS and added OptiMEM for imaging.

To record continuous 11.7 ms single-molecule movies, we stained the cells for 1 h with 10 nM HTL PA-JF-646. After incubation, we washed the cells two times with PBS and incubated them further in DMEM for 45 min at 37 °C and 5% CO_2_. Before imaging, we washed the cells three times with PBS and imaged them in OptiMEM.

### Single-molecule microscope setup

We conducted single-molecule imaging on a custom-built fluorescence microscope ^71^. It is built around a conventional Nikon body (TiE, Nikon) equipped with an AOTF (AOTFnC-400.650-TN, AA Optoelectronics), a high-NA objective (100x, NA 1.45, Nikon), a 638 nm laser (IBEAM-SMART-640-S, 150 mW, Toptica) and a 405 nm laser (Laser MLD, 299 mW, Solna, Sweden). For a good signal-to-noise ratio, we illuminated cells with a highly inclined and laminated optical sheet (HILO) ^36^. To detect the emitted fluorescence light, which previously passed a multiband emission filter (F72-866, AHF, Tübingen, Germany), we used an EMCCD camera (iXON Ultra DU 897, Andor, Belfast, UK).

### Single-molecule imaging

To assess the chromatin residence time of Halo-tagged molecules and to differentiate between unbinding of molecules and photobleaching, we performed time-lapse (tl) microscopy. Therefore, we recorded movies using a tl-cycle, which consisted of an image with a fixed camera integration time of 51.7 ms followed by a certain dark time. The tl-cycle times were 0.1 s, 0.4s, 1.6 s, and 6.4 s. Overall, movies covered 30 s, 120 s, 480 s, and 960 s, respectively. To avoid variances in the photobleaching rate, we adjusted the laser power to 1.13 mW before starting each measurement.

To track fast-moving molecules and finally determine diffusion coefficients and bound fractions, we recorded movies with a short exposure time of 10 ms, with a total frame cycle time of 11.7 ms. To avoid high background, we activated the fluorophore with 0.05 mW UV illumination for a short time period of 1 ms between two 638 nm exposures. Since we only activated a small fraction of fluorophores in every frame, it was feasible to record 20,000 frames without decreasing density of activated molecules.

### Single-molecule data analysis

We used TrackIT to identify, localize and track single molecules and perform diffusion analysis ^37^. For time-lapse imaging movies, we used the threshold factor 4.5 to identify spots. For connecting bound molecules through consecutive frames, we set the tracking radius to 0.9 pixels (0.1 s tl), 1.19 pixels (0.4 s tl), 1.75 pixels (1.6 s tl), and 2.8 pixels (6.4 s tl). The minimum tracking length was 3 frames for 0.1 s tl and 0.4 s tl and 2 frames for the other tl conditions. We allowed 2 gap frames without detection for 0.1 s tl and 1 gap frame for longer tl conditions. The gap frame was only allowed if the track already included 2 frames before the gap.

For 11.7 ms continuous movies, we detected spots with the threshold factor of 4. We connected tracks using a tracking radius of 7, a minimum track length of 2, gap frame of 1 and track length before gap frame of 2. To determine diffusion coefficients and fractions, we fitted the cumulative survival time distribution with a three-exponential Brownian diffusion model ^72^. The bin size of the distribution was set to 1120 which corresponded to 1 nm. For analysis we considered the first 5 jumps per track to avoid overrepresentation of immobile molecules and discarded jumps over gap frames. We estimated errors of diffusion coefficients *D*_*1,2,3*_ and fractions *A*_*1,2,3*_ by repeating diffusion analysis 500 times using 80% of randomly chosen jump distances. The amplitude *A*_*1*_ of the slowest diffusion component represented the overall bound fraction *f*_*b*_ of the tracked molecules. The unbound fraction of tracked molecules is then given by *p*_*f*_ *=1-f*_*b*_.

### Analysis of survival time distributions using GRID

We inferred dissociation rate spectra of HT-RBPJ variants by analyzing survival time distributions from time-lapse imaging with GRID ^38^. In brief, GRID reveals the amplitudes of *l* dissociation rates *k*_*off,l*_ associated with *l* binding classes from fluorescence survival time distributions by an inverse Laplace transformation. Initially, GRID reveals the event spectrum, whose amplitudes *A*^*e*^_*l*_ represent the relative frequency of binding events that occur for a certain binding class *l* over the observation period. *A*^*e*^_*l*_ is given by ^38^

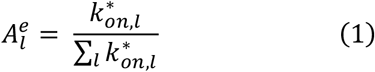

where *k\**_*on,l*_ *= k*_*on,l*_ *[D*_*l*_*], k*_*on,l*_ is the bimolecular association rate to a binding site of class *l*, and *[D*_*l*_] is the concentration of unoccupied binding sites *D*_*l*_ of type *l*. By dividing the amplitudes with the corresponding rates and renormalization, the event spectrum is converted to the state spectrum, with amplitudes *A*^*s*^_*l*_^38^:

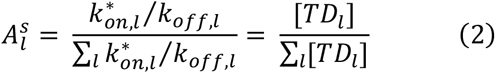

where *[TD*_*l*_] is the concentration of transcription factors *T* bound to binding sites *D*_*l*_ of class *l*. The amplitudes *A*^*s*^_*l*_ of the state spectrum reflect the probability to find a molecule in a certain binding class *l* at any time snapshot. Together with the overall bound fraction *f*_*b*_, we obtain the overall fraction of molecules binding to a binding site of class *l*: *p*_*b,l*_ = *f*_*b*_ * *A*^*s*^_*l*_.

We estimated the error of dissociation rate spectra by repeating the GRID analysis 499 times with 80% of randomly chosen survival times for each GRID run.

### Calculation of the target site search time for the search mechanism of facilitated diffusion

In a commonly assumed model of transcription factor – DNA interactions, the transcription factor has two different classes of binding sites on DNA, unspecific and specific ^1,39^. It thus performs the unspecific and specific binding reactions:

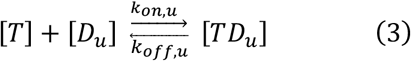

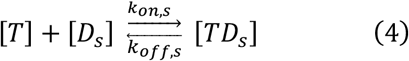

Where *[T]* is the concentration of the unbound protein, *[D*_*u*_] and *[D*_*s*_] are the concentrations of unoccupied unspecific and specific binding sites, *[TD*_*u*_] and *[TD*_*s*_] are the concentrations of protein-bound binding sites, *k*_*on,u*_ and *k*_*on,s*_ are the bimolecular association rates and *k*_*off,u*_ and *k*_*off,s*_ are the dissociation rates (see analysis of survival time distributions).

The dissociation constant of either reaction is given by

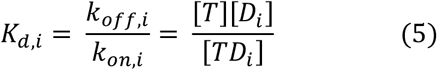

with *i = u,s*. Thus, there is a relation between the ratio of kinetic rates and the ratio of the unbound fraction *p*_*f*_ (see single-molecule data analysis) to the bound fraction *p*_*b,i*_ (see analysis of survival time distributions) of the DNA-binding protein:

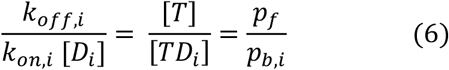

We set *k\**_*on,i*_ *= k*_*on,i*_ *[D*_*i*_] in the following.

To search for their specific target sites, transcription factors follow a search mechanism of facilitated diffusion including three-dimensional diffusion in the nucleoplasm and one-dimensional sliding along unspecific DNA ^39^. We previously devised a three-state model of the target site search mechanism of facilitated diffusion (Figure 3a) ^43^. Within this model, the transcription factor may occupy the three states: unbound (or free), unspecifically bound, and specifically bound. Unspecific binding with association rate *k\**_*on,u*_ *= k*_*on,u*_ *[D*_*u*_] and dissociation rate *k*_*off,u*_ may occur both at unspecific binding sites and in the vicinity of a specific target site. Specific binding may be achieved by either transition of the transcription factor from unspecific binding to specific binding with the microscopic on-rate *k*_*u-s*_ while sliding over the specific target site, or by direct association with the association rate *k\**_*on,s*_ = *k*_*on,s*_ *[D*_*s*_]. These two pathways are combined in the effective specific association rate *k\**_*a*_. Dissociation from a specific binding site may occur either by transition to the unspecifically bound state with the microscopic off-rate *k*_*s-u*_, or by direct dissociation with dissociation rate *k*_*off,s*_. These two pathways are combined in the effective specific dissociation rate *k*_*d*_ = 1 / τ_*s*_. The target site search time τ_*search*_ = *1 / k\**_*a*_ can be found as the inverse of the effective specific association rate. It is the average time a transcription factor needs to find any of the specific target sites and includes multiple rounds of unspecific binding and unbinding at unspecific binding sites or in the vicinity of the specific target site, before a final transition to specific binding occurs.

The association and dissociation rates of the three-state model are coupled by detailed balance^43^:

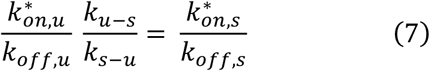

Of the parameters entering the three-state model of facilitated diffusion, the unspecific dissociation rate *k*_*off,u*_ and the effective specific dissociation rate *k*_*d*_ are experimentally accessible. They are found by GRID analysis of the fluorescence survival time distributions of the transcription factor binding times measured with time-lapse microscopy (see analysis of survival time distributions). Moreover, the fractions of unbound (*p*_*f*_), unspecifically bound (*p*_*u*_) and specifically bound (*p*_*s*_) molecules are experimentally accessible and found by diffusion measurements and GRID analysis (see analysis of single-molecule data and analysis of survival time distributions).

The unspecific association rate *k\**_*on,u*_ can be obtained by:

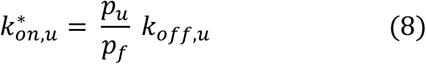

To compare the search times of different HT-RBPJ variants despite we did not have access to the rates *k*_*u-s*_, *k*_*s-u*_, *k\**_*on,s*_, and *k*_*off,s*_, we fixed the ratio *k*_*off,s*_ / *k*_*d*_ for all variants to 0.0705, corresponding to a target site search time ten times faster than direct specific association. The direct specific association rate *k\**_*on,s*_ can be obtained with equation 6:

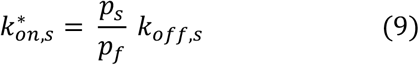

In the three-state model, the direct specific dissociation rate *k*_*off,s*_ is coupled to *k\**_*on,u*_, *k*_*u-s*_, and *k*_*d*_ by^43^:

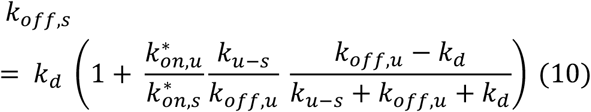

Thus, for given *k\**_*on,u*_, *k*_*d*_, and *k*_*off,s*_, the microscopic on-rate *k*_*u-s*_ can be obtained:

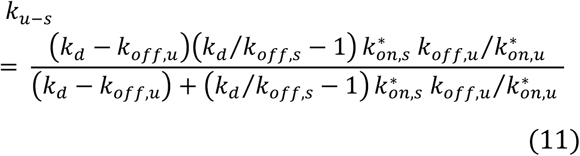

and the microscopic off-rate *k*_*s-u*_ is found from equation 11 and equation 7 of detailed balance:

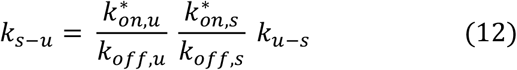

Finally, the target site search time τ_*search*_ of the three-state model is given by ^43^:

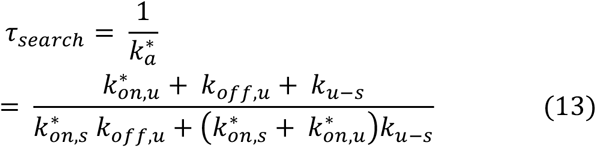

The parameters entering the model and the resulting target site search times can be found in Supplementary Table 7.

To obtain τ_*search*_ for a given direct specific association rate *k\**_*on,s*_, we solved equation 9 for *k*_*off,s*_ and followed the subsequent steps to calculate the target site search time.

### Construction of the binding energy landscape

We determined the binding energy landscape of a transcription factor from the experimentally accessible kinetic rates *k*_*off,u*_ and *k*_*d*_, the measured unbound and bound fractions *p*_*f*_, *p*_*u*_, and *p*_*s*_, and the kinetic rates *k\**_*on,u*_, *k*_*u-s*_, *k*_*s-u*_, and *k\**_*on,s*_ obtained from the three-state model of facilitated diffusion. Importantly, while binding energies are often compared at standard concentrations of binding sites, we judged it more relevant to compare the binding energies of transcription factor variants at the respective concentration of binding sites they actually have in the nucleus. The binding energy Δ*G*_*u*_ of the unspecifically bound state is given by:

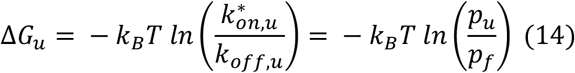

with the Boltzmann constant *k*_*B*_ and the unit of thermal energy *k*_*B*_*T*. Analogously, the binding energy Δ*G*_*s*_ of the specifically bound state is given by:

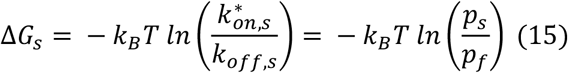

To compare the relative height of energy barriers of the various association and dissociation transitions, we assumed a similar frequency factor *k*_*A*_ for all transitions. The energy barriers of unspecific binding (Δ*G*_*f*→*u*_), of switching from unspecific to specific binding (Δ*G*_*u*→*s*_), and of specific binding (Δ*G*_*f*→*s*_) are then given by:

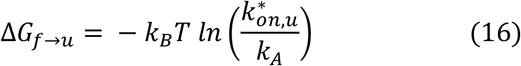

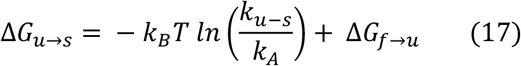

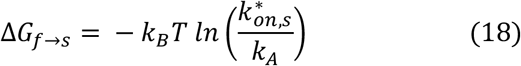

## Acknowledgements

We thank Jutta Hegler for help with Lentivirus production and Sabine Schirmer for excellent technical assistance, Devin Assenheimer for help with error calculations and Jonas Coßmann and Tobias Bischof for supportive discussions (all Ulm University). Halo-SiR ligand was kindly provided by K. Johnsson (Max Planck Institute for Medical Research). pLV-tetO-Oct4 was a gift from Konrad Hochedlinger (Massachusetts General Hospital), pMD2.G and psPAX2 were a gift from Didier Trono (Ecole Polytechnique Fédérale de Lausanne). pSpCas9(BB)-2A-Puro (PX459) V2.0 was a gift from Feng Zhang. We thank the Core Facility FACS of Ulm University for their help with cell sorting, with special thanks to Dr Simona Ursu, Dr. Sarah Warth, and Daniela Froelich.

The work was funded by the Deutsche Forschungsgemeinschaft (DFG, German Research Foundation no. 427512076 to J.C.M.G. and F.O., no. 468578170 and 422780363 SPP 2202 to J.C.M.G.), the European Research Council (ERC) under the European Union’s Horizon 2020 Research and Innovation Program (no. 637987 ChromArch to J.C.M.G.), and the German Cancer Aid (#70114289 to F.O.) R.K. acknowledges funding by NSF/MCBBSF grant #1715822, T.B. acknowledges funding by the DFG TRR81-A12, a research grant of the University Medical Center Gießen and Marburg (UKGM), the LOEWE research cluster iCANx, and the Excellence Cluster for Cardio Pulmonary System (ECCPS) in Gießen. M.B. acknowledges funding by the Forschungscampus Mittelhessen. Support by the Collaborative Research Centres 1074 (DFG no 217328187), 1279 (DFG no. 316249678) and 1506 (DFG no. 450627322) and the Center for Translational Imaging MoMAN of Ulm University (DFG no. 447235146) is acknowledged.

## Author contributions

F.O. and J.C.M.G. conceived the project; D.H., P.H., F.O. and J.C.M.G. designed the project; F.F and B.D.G. generated the knockout cell lines; P.H. and F.O. performed luciferase assays; F.F., B.D.G. and T.B. performed ChIP-Seq experiments; T.F., B.D.G. and M.B. analyzed ChIP-Seq data; R.K. analyzed the RBPJ structure; D.H. performed the single-molecule measurements; D.H. and J.C.M.G. analyzed the single-molecule data with contributions from K.Z.; D.H., P.H., F.O. and J.C.M.G wrote the manuscript with comments from all authors.

## Data availability

Source data for figures are provided as a separate file named ‘Source Data.xlsx’. ChIP-Seq data was uploaded to the Gene Expression Omnibus repository (GEO accession number: GSE249973, reviewer-token: ibarkqiovdkprqd). Single-particle tracking data are available at Data Dryad repository and can be accessed during review with the link https://datadryad.org/stash/share/H5P9V3BYcN3Hxi8f_6xWl42yKexKgjHgPjSvRCZ1DLU. It will be freely available after publication. Data supporting the findings of this manuscript will be available from the corresponding authors upon reasonable request.

## Code availability

The single-molecule tracking software TrackIt is freely available on GitLab (https://gitlab.com/GebhardtLab/TrackIt).

