## Supplementary Information for "Effective *in vivo* binding energy landscape illustrates kinetic stability of RBPJ-DNA binding"

Supplementary Figures

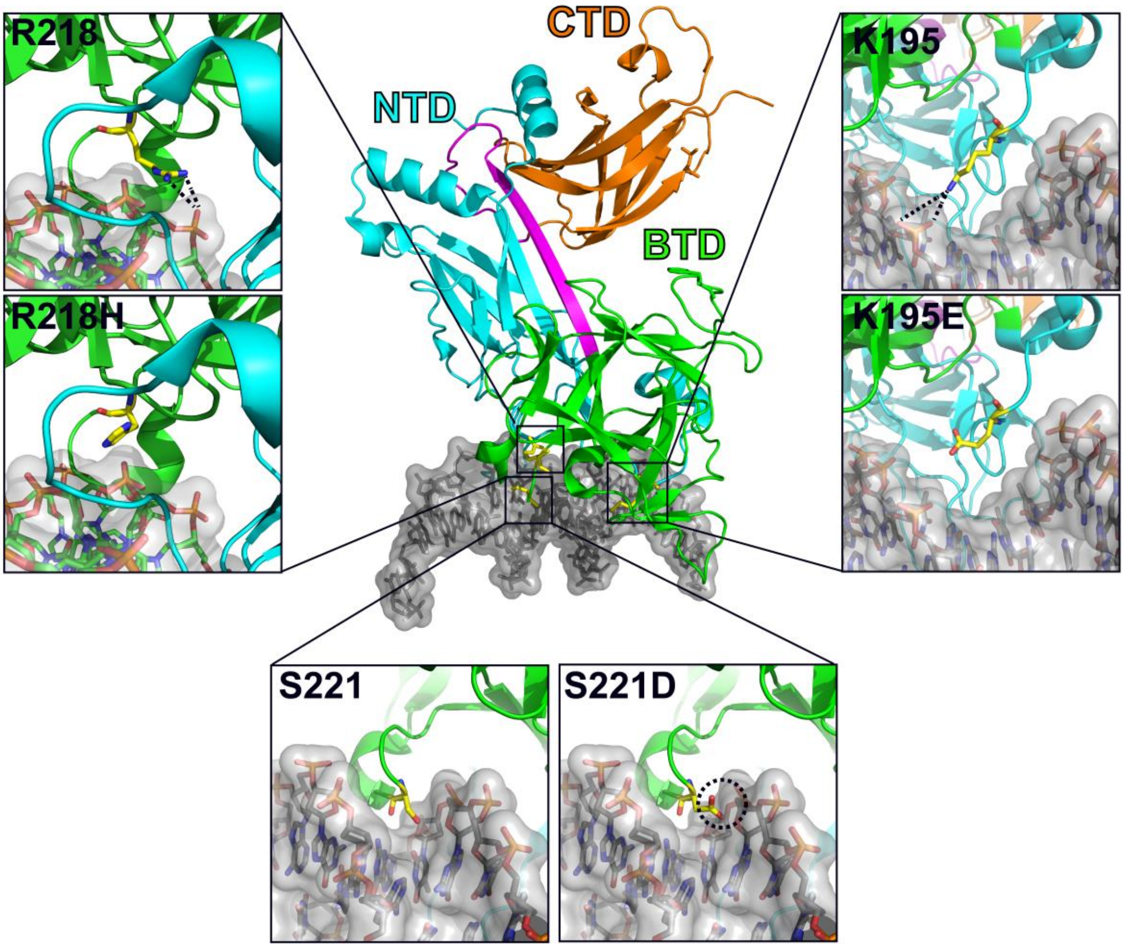

**Supplementary Figure 1: Spatial orientation of DNA binding interface mutations of RBPJ**

Structure of mouse RBPJ in complex with the HES1 non-consensus site (PDB entry: 3IAG) with highlighted NTD (cyan), BTD (green) and CTD (orange) as well as the DNA (grey). Zooms: Spatial orientation of the amino acids K195, R218 and S221 and of the DNA-binding deficient mutations K195E, R218H and S221D. Structures and mutations of single amino acids were performed using PyMOL.

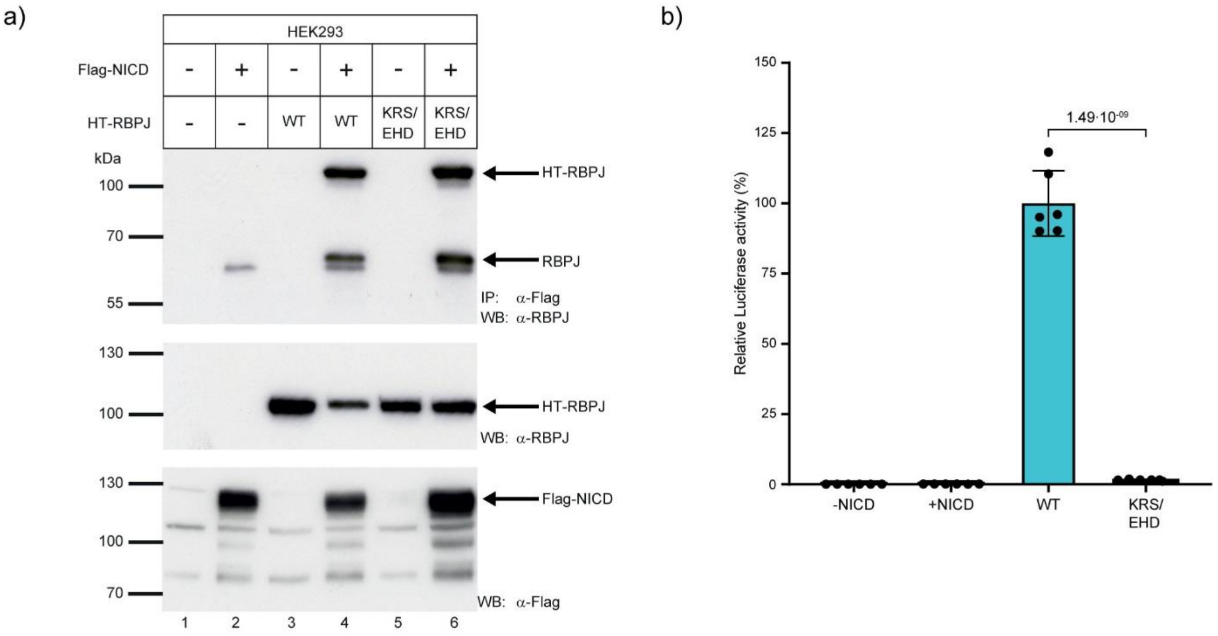

**Supplementary Figure 2: Controls for HT-tagged RBPJ variants**

**a)** HT-RBPJ-WT and HT-RBPJ-KRS/EHD show comparable interaction with Flag-NICD. HEK293 cells were co-transfected with expression plasmids for Flag-tagged NICD and Halo-tagged RBPJ-WT or -KRS/EHD. Co-IP was performed 24 h after transfection using an anti-Flag antibody (upper panel). Western blotting was performed to verify expression of Halo-tagged RBPJ proteins (middle panel) and Flag-tagged NICD (bottom panel). **b)** In vivo Luciferase activity assays. Relative luciferase activity of RBPJ-specific reporter constructs in RBPJ knockout cell line (#42) transfected with HT-RBPJ-WT or -KRS/EHD and co-transfected with NICD (mean values ± s.d.). P-values were determined using two-tailed, unpaired Student's t-test. -/+NICD: control with untransfected / NICD-transfected RBPJ knockout cell line. For unprocessed Western blots see Source Data file.

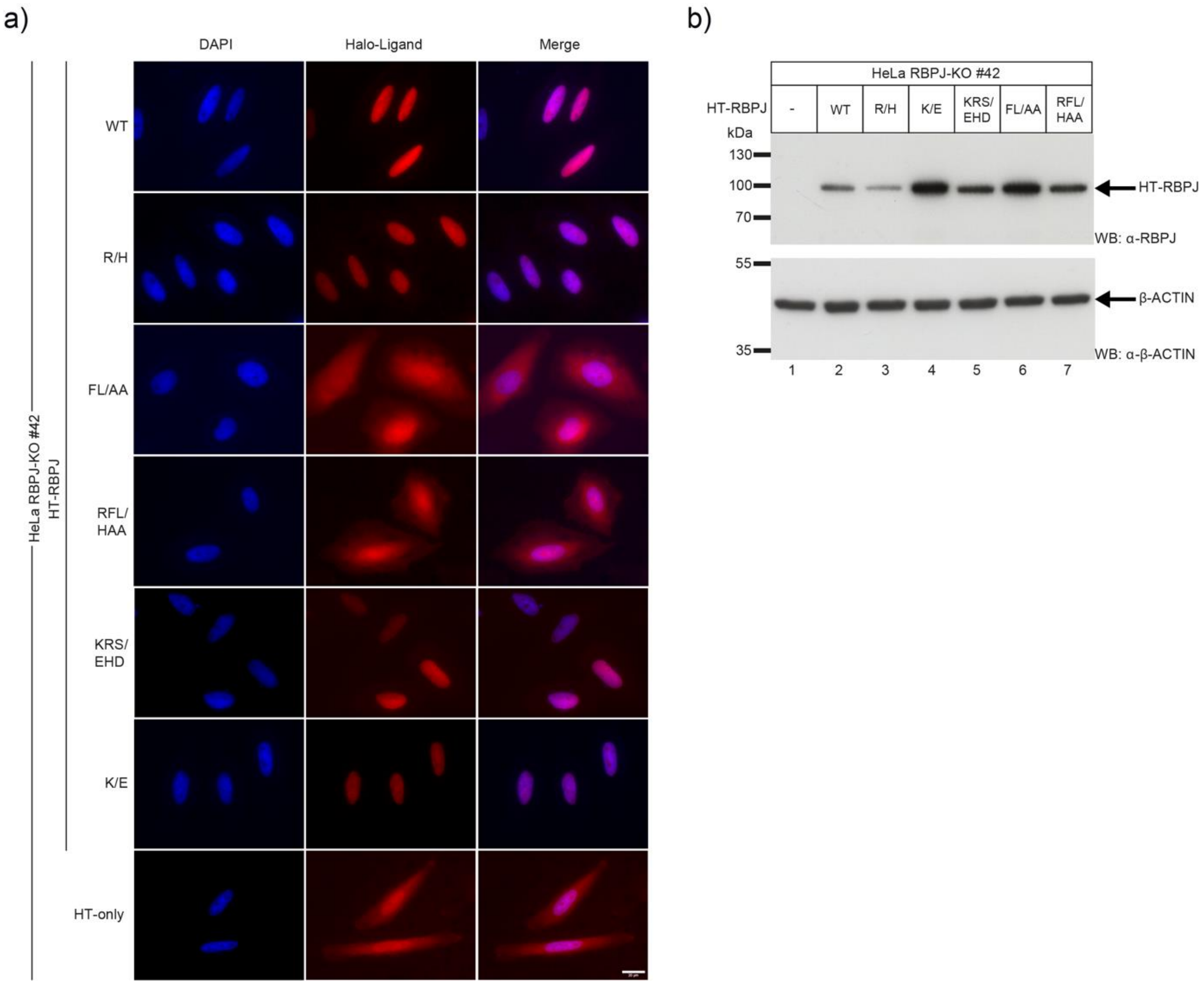

**Supplementary Figure 3: Characterization of RBPJ-depleted HeLa cell lines with reintroduced RBPJ variants**

**a)** Fluorescence images of RBPJ-depleted HeLa cells (clone #42) stably expressing HT-RBPJ-WT, HT-RBPJ variants or the HaloTag alone. Nuclei were labeled with DAPI (Column 1) and HaloTag was fluorescently labeled with HTL-TMR (Column 2). Merged images (Column 3) revealed nuclear distribution of HT-RBPJ-WT and RBPJ variants with mutations in the DNA binding interface only (R/H, KRS/EHD and K/E). RBPJ variants with mutations in the cofactor binding interface (FL/AA and RFL/HAA) as well as HaloTag alone showed both cytoplasmic and nuclear distribution. Images were taken with a fluorescence microscope with a 63x lens and the scale bar represents 20 µm. For abbreviation of mutations see main Text.

**b)** Verification of stable expression of HT-RBPJ-WT and variants in RBPJ-depleted HeLa cell line #42 using Western blot. Lanes 2-7 contained lysates from cell lines expressing the following RBPJ variants, respectively: WT, R/H, K/E, KRS/EHD, FL/AA and RFL/HAA. Lane 1 contained lysate of RBPJ-depleted cells and was used as negative control. A specific $\alpha$-RBPJ antibody was used for blotting. Detection of $\beta$-ACTIN was used as loading control using a specific $\alpha$-$\beta$-ACTIN antibody. Arrows indicate bands for HaloTag-RBPJ or $\beta$-ACTIN. For unprocessed Western blot see Source Data file. For abbreviation of mutations see main Text.

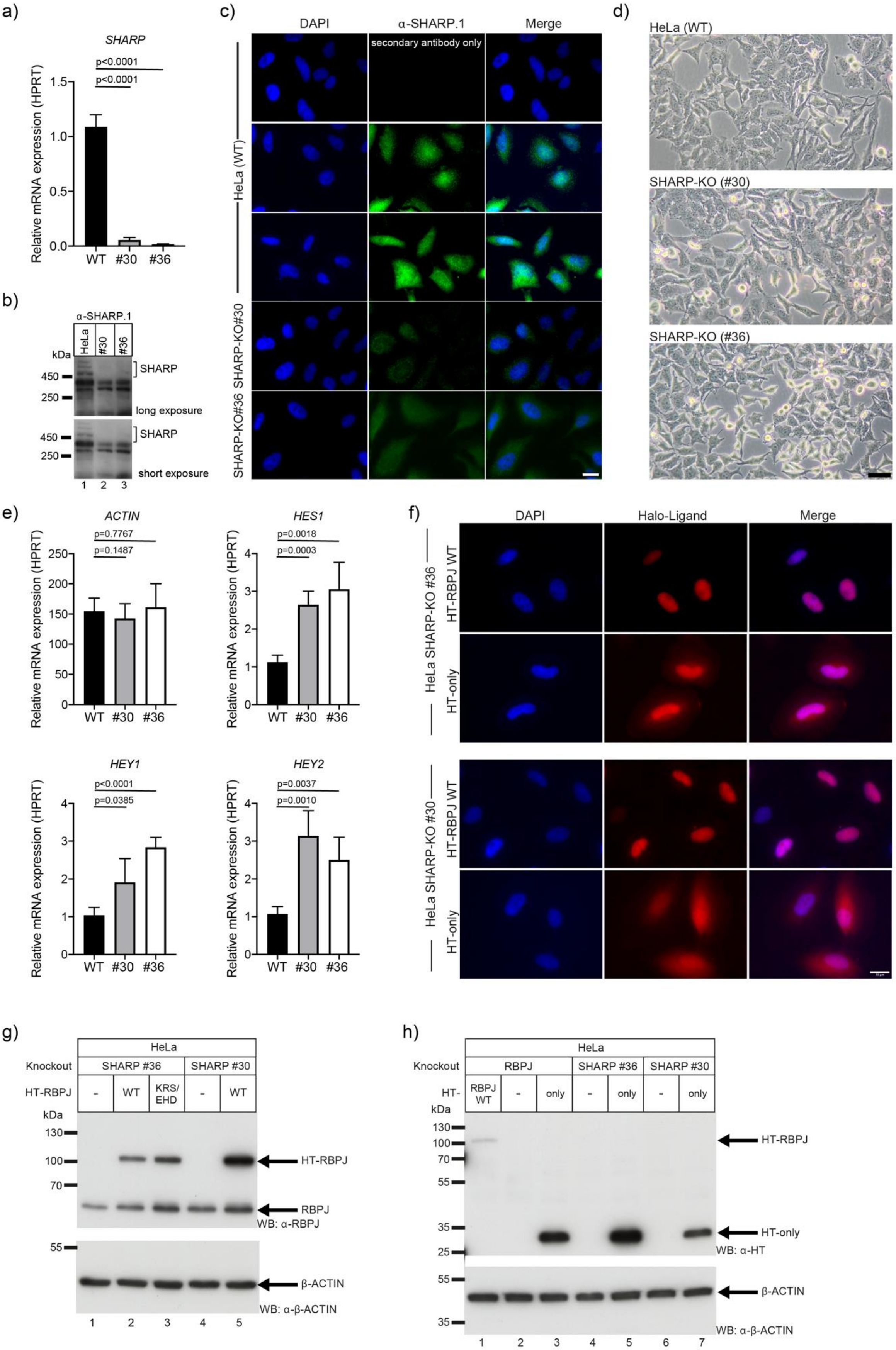
a)
SHARP
p<0.0001
p<0.0001
Relative mRNA expression (HPRT)
WT
#30
#36
b)
α-SHARP.1
HeLa
#30
#36
kDa
SHARP
450
250
long exposure
short exposure
c)
DAPI
α-SHARP.1
Merge
secondary antibody only
HeLa (WT)
SHARP-KO#30
SHARP-KO#36
d)
HeLa (WT)
SHARP-KO (#30)
SHARP-KO (#36)
e)
ACTIN
p=0.7767
p=0.1487
HES1
p=0.0018
p=0.0003
HEY1
p<0.0001
p=0.0385
HEY2
p=0.0037
p=0.0010
f)
DAPI
Halo-Ligand
Merge
HeLa SHARP-KO #36
HeLa SHARP-KO #30
HT-RBPJ WT
HT-only
g)
HeLa
Knockout
SHARP #36
SHARP #30
HT-RBPJ
WT
KRS/ EHD
HT-RBPJ
RBPJ
WB: α-RBPJ
β-ACTIN
WB: α-β-ACTIN
h)
RBPJ
only
RBPJ WT
HT-only
WB: α-HT

**Supplementary Figure 4: Characterization of SHARP-depleted HeLa cell lines**

**a)** Relative expression of SHARP mRNA in control and SHARP-depleted HeLa cell lines (clones #30 and #36). Expression was determined by qPCR and calculated relative to the housekeeping gene HPRT. SHARP-depleted cells display almost full depletion of SHARP expression. Shown is the mean value ± s.d. originating from 4 independent experiments, each containing at least 1 technical replicate. P-values were determined using two-tailed, unpaired Student's t-test.

**b)** Detection of SHARP in control and SHARP-depleted HeLa cells by Western blotting. Two distinct bands were detected at the height of ~500 kDa in HeLa control cells by using an α-SHARP.1 antibody. Long- and short-term exposures are displayed in the upper and lower panel, respectively.

**c)** Immunofluorescence microscopy images using the α-SHARP.1 antibody. Images were taken with a 63x lens and the scale bar represents 20 µm. Control and SHARP-depleted HeLa cells #30 and #36 were stained with DAPI for visualization of the nucleus (Column 1). As negative control, HeLa control cells were incubated only with secondary antibodies (row 1), while the other samples were additionally incubated with a primary α-SHARP.1 antibody (row 2 to 5). SHARP-specific fluorescence signals are shown in column 2. Merged images (Column 3) show the subcellular localization of SHARP in the nucleus and in the cytoplasm in all cell lines, however, SHARP-depleted cells #30 and #36 show a reduced fluorescent signal (Row 4 and 5).

**d)** Bright-field microscopy images of control (top panel) and SHARP-depleted HeLa cells #30 (middle panel) and #36 (bottom panel). The images were taken using a 20x lens and the scale bar represents 50 µm.

**e)** mRNA expression levels of Notch target genes *HES1*, *HEY1* and *HEY2* as well as of the housekeeping gene *ACTIN* were determined by qPCR. HeLa control and HeLa SHARP-depleted cells #30 and #36 were examined. Except for the housekeeping gene *ACTIN*, SHARP-depleted cells exhibited increased mRNA levels of Notch target genes compared to HeLa control cells due to derepression. Shown are the means ± s.d. originating from 4 independent experiments, each containing at least 1 technical replicate. P-values were determined using two-tailed, unpaired Student's t-test.

**f)** Fluorescence microscopy images of SHARP-depleted HeLa cell lines #36 and #30 stably expressing HT-RBPJ-WT or just the HaloTag. The nuclei were stained with DAPI (Column 1), HaloTag was labeled with HTL-TMR (Column 2). Merged images (Column 3) show that HT-RBPJ-WT is located exclusively in the nucleus, HaloTag alone is located in both the cytoplasm and the nucleus.

**g)** Verification of stable expression of HT-RBPJ-WT and HT-RBPJ-KRS/EHD using Western blotting. Lanes 2 and 3 contained lysates from SHARP-depleted HeLa cells #36 expressing either HT-RBPJ-WT or HT-RBPJ-KRS/EHD. Lane 5 contained lysate from SHARP-depleted HeLa cells #30 expressing HT-RBPK-WT. Lane 1 and 4 contained lysates of SHARP-depleted cells #36 and #30 as negative controls. A specific α-RBPJ antibody was used. Detection of β-ACTIN was used as loading control using a specific α-β-ACTIN antibody. Arrows indicate bands for endogenous RBPJ, HT-RBPJ-WT or β-ACTIN. Unprocessed Western Blots are provided as Source Data file.

**h)** Verification of stable expression of HT-RBPJ-WT or HaloTag protein using Western blotting. Lanes 3, 5 and 7 contained lysates from RBPJ-depleted, SHARP-depleted #36 or SHARP-depleted #30 cell lines expressing the HaloTag protein. Lanes 2, 4 and 6 contained lysates from RBPJ-depleted, SHARP-depleted #36 or SHARP-depleted #30 cell lines respectively as negative controls. Lane 1 contained lysate from a RBPJ-depleted cell line expressing HT-RBPJ-WT as a positive control. A specific $\alpha$-HaloTag antibody was used. Detection of $\beta$-ACTIN was used as loading control using a specific $\alpha$-$\beta$-ACTIN antibody. Arrows indicate bands for HaloTag protein, HT-RBPJ-WT or $\beta$-ACTIN. For unprocessed Western Blot see Source Data file.

Source data are provided as Source Data file for Supplementary Figure 4a,e.

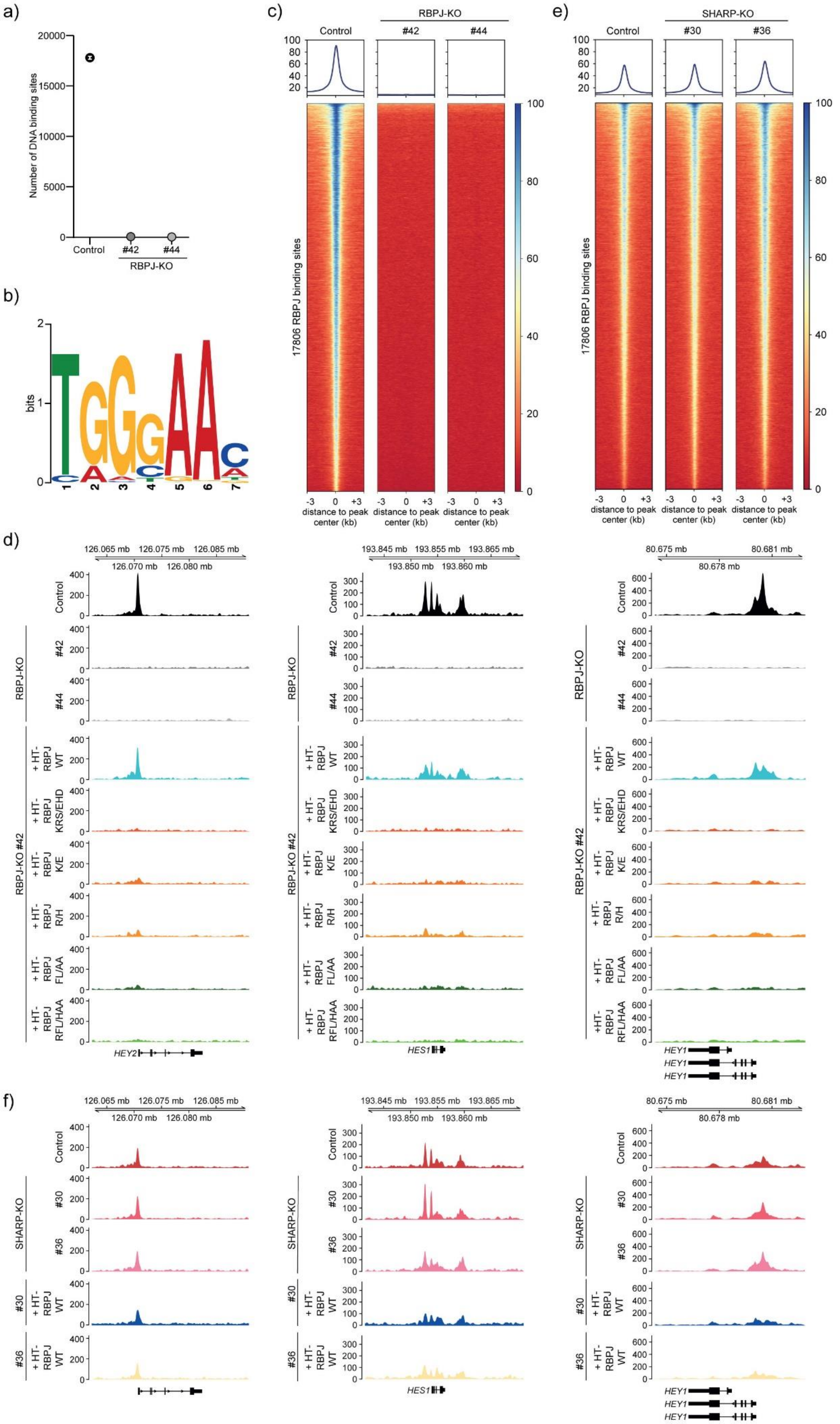
a)
Number of DNA binding sites
Control
#42
#44
RBPJ-KO
b)
bits
c)
Control
RBPJ-KO
#42
#44
17806 RBPJ binding sites
distance to peak center (kb)
e)
SHARP-KO
#30
#36
d)
HEY2
HES1
HEY1
+ HT-RBPJ WT
+ HT-RBPJ KRS/EHD
+ HT-RBPJ K/E
+ HT-RBPJ R/H
+ HT-RBPJ FL/AA
+ HT-RBPJ RFL/HAA
RBPJ-KO #42
f)

**Supplementary Figure 5: ChIP-Seq analysis of RBPJ-depleted and SHARP-depleted cell lines**

**a)** Number of DNA binding sites identified in RBPJ ChIP-Seq for control HeLa cells transfected with an empty vector and RBPJ-depleted HeLa cell lines #42 and #44. In contrast to control HeLa cells, the RBPJ-depleted cell lines showed absence of RBPJ binding events. Error bars denote square root of the value.

**b)** RBPJ-DNA binding motif obtained from 17806 RBPJ occupied sites identified in HeLa control cells.

**c)** Heatmap of RBPJ ChIP-Seq reads covering 3 kb up- and downstream of binding sites for all 17806 binding sites identified in control HeLa cells transfected with an empty vector. In contrast to the control HeLa cells (Left column), the RBPJ-depleted cell lines #42 (Middle column) and #44 (Right column) showed absence of RBPJ binding events.

**d)** Genome browser sections of the location indicated on top showing the ChIP-Seq coverage for promoter or enhancer regions of the Notch target genes *HEY2* (left panel), *HES1* (middle panel), and *HEY1* (right panel) for RBPJ-depleted cells #42 and #44 and for RBPJ-depleted cells stably expressing the indicated HT-RBPJ variants.

**e)** Heatmap of RBPJ ChIP-Seq reads covering 3 kb up- and downstream of binding sites for all 17806 binding sites identified in control HeLa cells transfected with an empty vector. Compared to the control cell line (Left column), the two SHARP-depleted cell lines #30 (Middle column) and #36 (Right column) showed a similar profile of RBPJ binding events.

**f)** Genome browser sections of the location indicated on top showing the ChIP-Seq coverage for promoter or enhancer regions of the Notch target genes *HEY2* (left panel), *HES1* (middle panel), and *HEY1* (right panel) for the SHARP-depleted cells #30 and #36 and for SHARP-depleted cells stably expressing HT-RBPJ-WT.

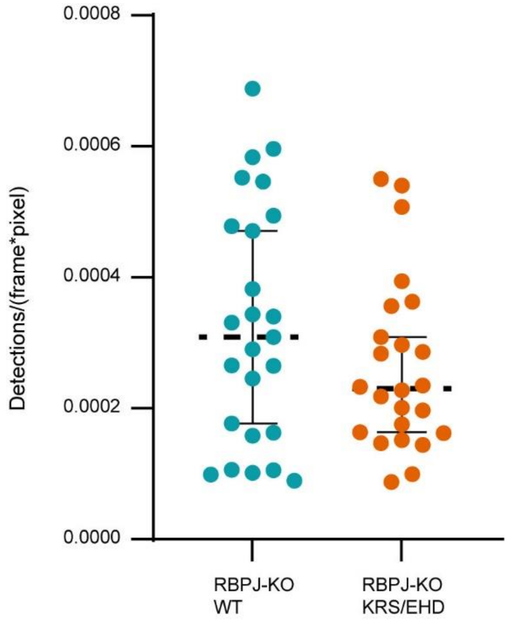

**Supplementary Figure 6:**

Average number of detected molecules per frame of 11.7 ms continuous movies for HT-RBPJ-WT and -KRS/EHD in RBPJ knockout cells (#42). Dashed black lines indicate the median value and the solid lines the 0.25 and 0.75 quartiles. Experimental statistics are listed in Supplementary Table 5. Data are provided as Source Data file for Supplementary Figure 6.

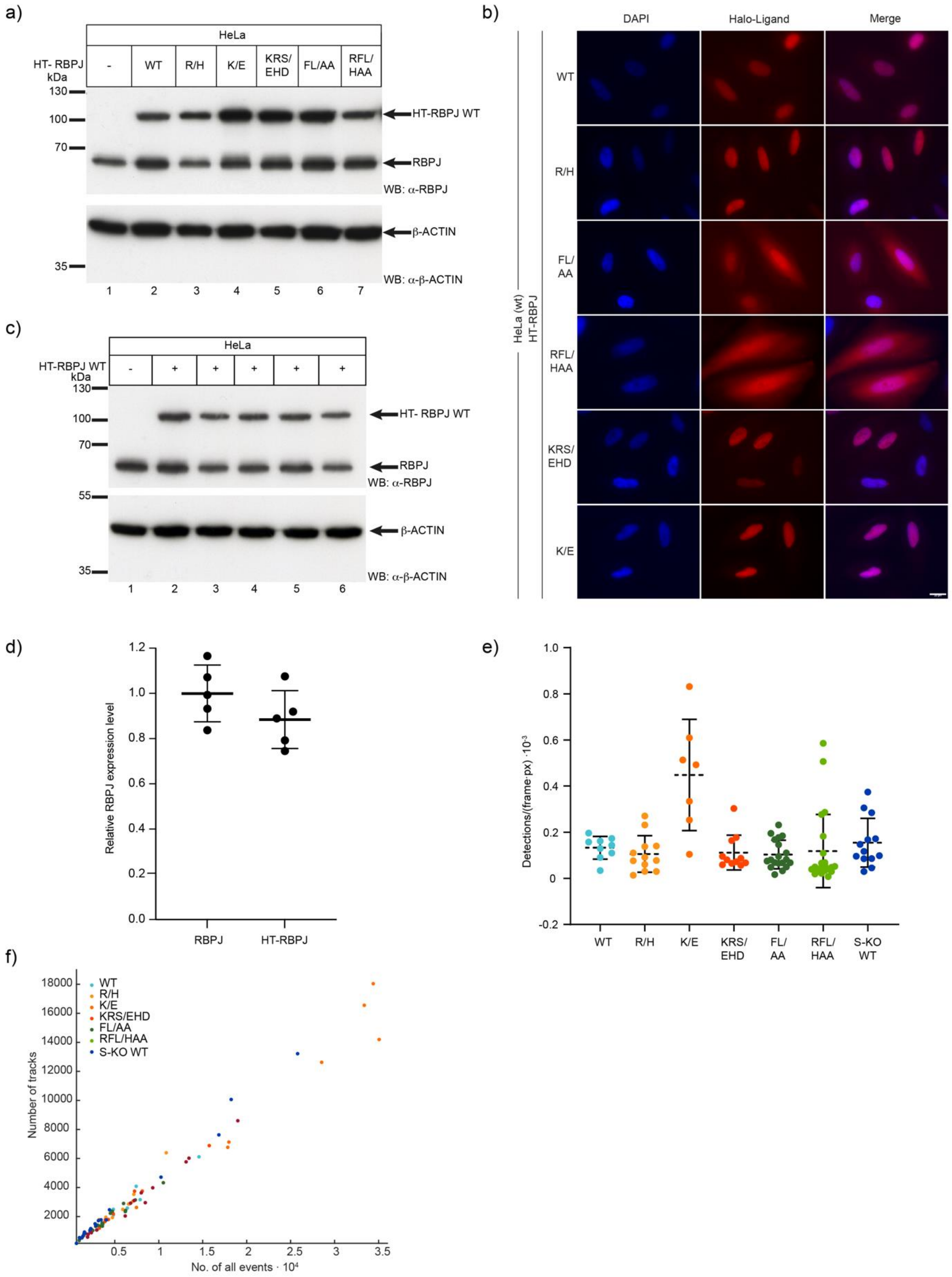

**Supplementary Figure 7: Quantification of expression level of HT-RBPJ variants**

**a)** Verification of stable expression of HT-RBPJ variants using Western blots. Lanes 2-7 contained lysates from HeLa cell lines expressing the following HT-RBPJ proteins respectively: wildtype, R218H, K195E, K195E/R218H/S221D, F261A/L388A and R218H/F261A/L388A. Lane 1 contained lysate of untreated HeLa cells as negative control. A specific α-RBPJ antibody was used. Detection of β-Actin was used as loading control using a specific α-β-Actin antibody. Arrows indicate bands of endogenous RBPJ, HT-RBPJ or β-Actin. For unprocessed Western blots see Source Data.

**b)** Fluorescence images of HeLa cells stably expressing HT-RBPJ variants. The nuclei were labeled with DAPI (Column 1) and HaloTag was fluorescently labeled with HTL-TMR (Column 2). Merged images (Column 3) revealed that HT-RBPJ-WT and mutants of the DNA binding interphase (R/H, KRS/EHD, and K/E) were localized exclusively in the nucleus (row 1, 2, 5 and 6). HT-RBPJ mutants FL/AA and RFL/HAA were localized in both the cytoplasm and the nucleus (row 3 and 4). Images were taken with a fluorescence microscope with a 63x lens and the scale bar represents 20 µm.

**c)** Verification of expression of endogenous RBPJ and HT-RBPJ-WT using Western blotting. Lanes 2 to 6 contained lysates of HeLa cell lines expressing HT-RBPJ-WT. Lane 1 contained lysate of HeLa cells as negative control. A specific α-RBPJ antibody was used. Detection of β-Actin was used as loading control using a specific α-β-Actin antibody. Arrows indicate bands of endogenous RBPJ, HT-RBPJ-WT or β-Actin. For unprocessed Western blots see Source Data.

**d)** Quantification of HT-RBPJ-WT expression level in HeLa cells using five independent lysates of HeLa cells stably expressing HT-RBPJ-WT. Results show mean relative RBPJ expression levels with standard deviation. In the Western blot of c), each band of endogenous RBPJ or HT-RBPJ-WT was marked with a region of interest of similar size and the mean intensity of each band was determined. The mean intensity of all endogenous RBPJ bands was determined, normalized to 1 and used as a base level for RBPJ expression. The individual expression levels of HT-RBPJ-WT were calculated by dividing the intensity of each individual HT-RBPJ-WT band by the mean intensity of all endogenous RBPJ bands to determine the relative intensity values. The final expression level for HT-RBPJ-WT was determined by calculating the mean of all individual relative intensity values.

**e)** Average number of detected single-molecule spots per frame for indicated HT-RBPJ variants. Dashed black lines indicate the median value and the solid lines the 0.25 and 0.75 quantiles. Experimental statistics are listed in Supplementary Table 5.

**f)** Number of identified single-molecule tracks versus the number of detected spots per frame for indicated HT-RBPJ variants. Data shown are from 10 ms continuous movies. Experimental statistics are listed in Supplementary Table 5.

Source data of this graph are provided as Source Data file for Supplementary Figure 7d,e,f.

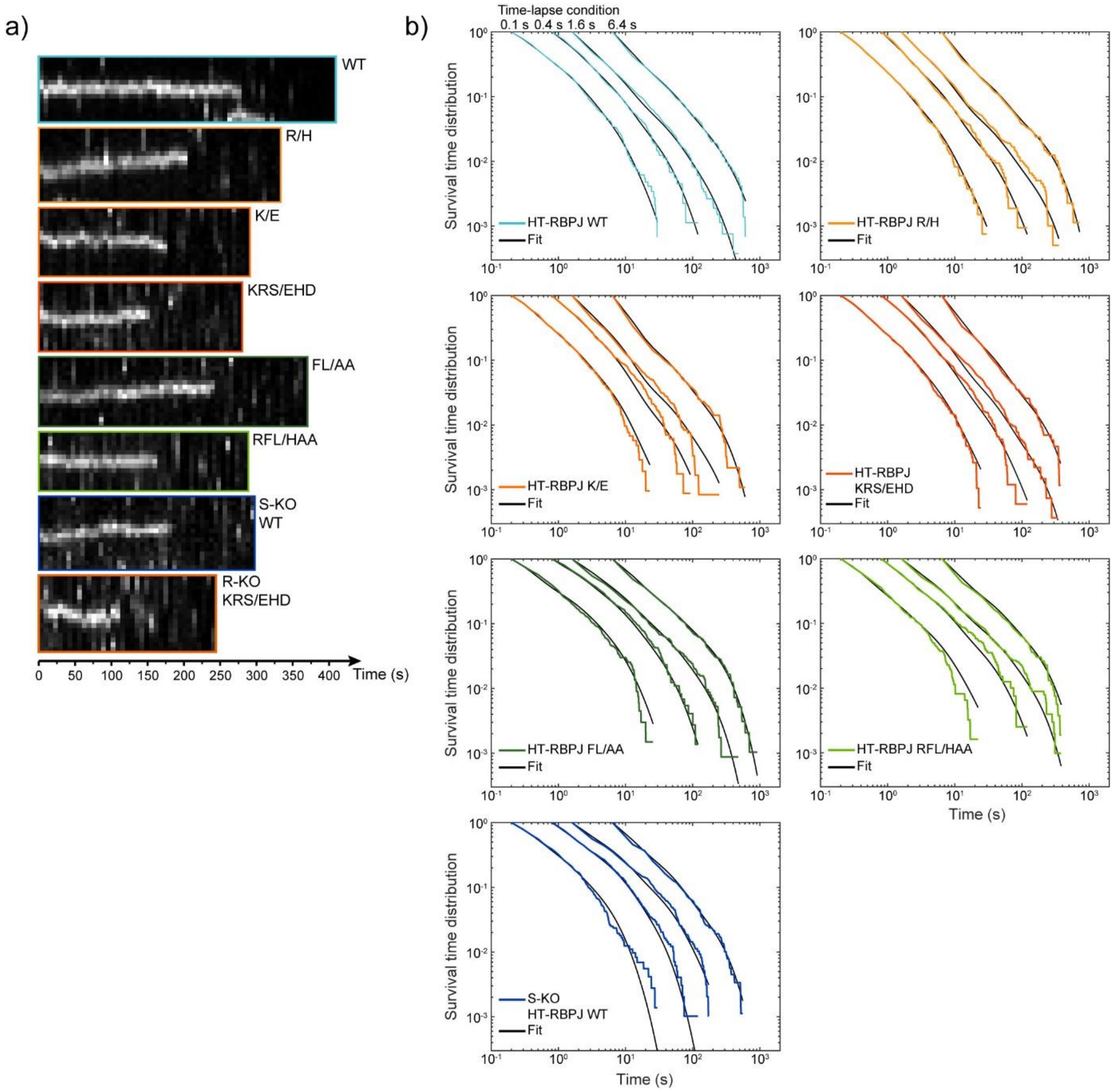

**Supplementary Figure 8: Survival time distributions of HT-RBPJ variants**

**a)** Example kymographs of a bound HT-RBPJ molecule of the indicated variant from 6.4 s time-lapse measurements.

**b)** Survival time distributions HT-RBPJ variants at time-lapse conditions shown on top and survival time function obtained with GRID (black lines). Experimental statistics are listed in Supplementary Table 3.

Source data are provided as Source Data file for Supplementary Figure 8b.

a)

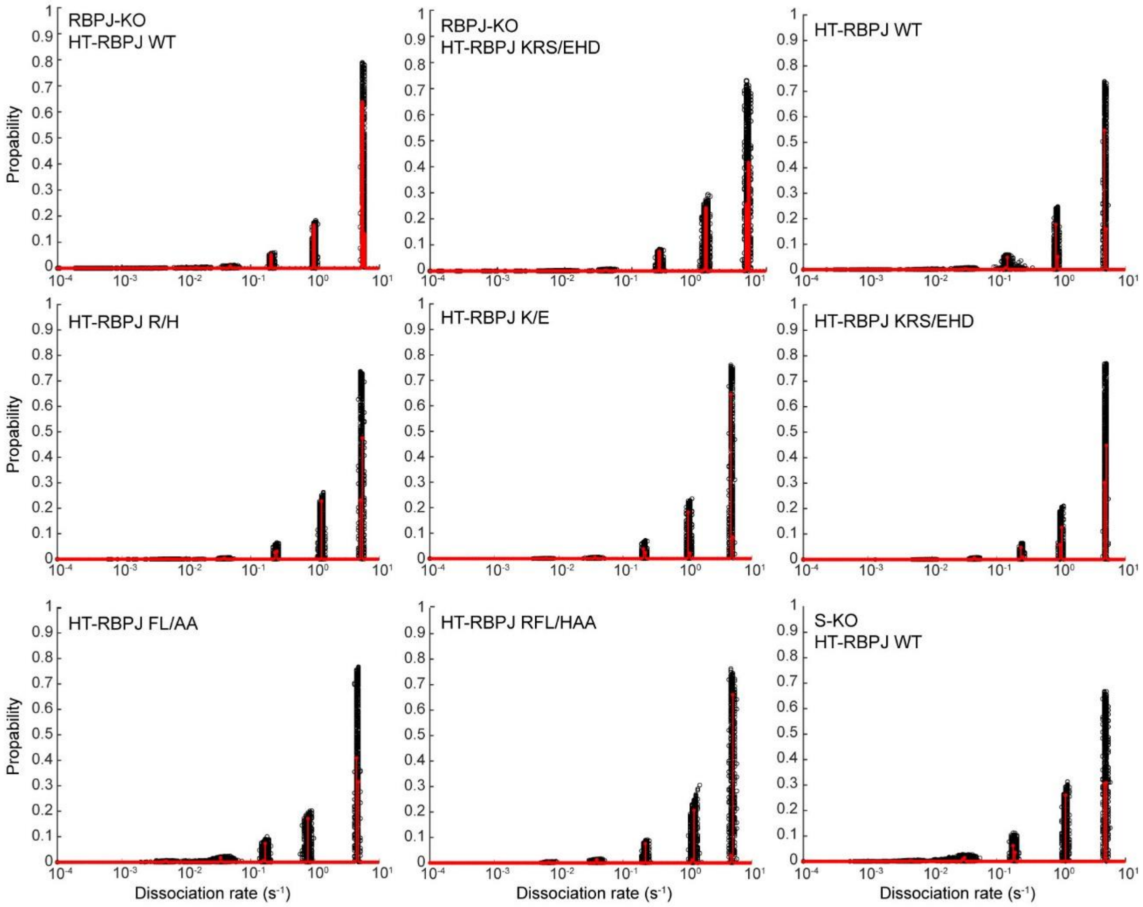
RBPJ-KO
HT-RBPJ WT
RBPJ-KO
HT-RBPJ KRS/EHD
HT-RBPJ WT
HT-RBPJ R/H
HT-RBPJ K/E
HT-RBPJ KRS/EHD
HT-RBPJ FL/AA
HT-RBPJ RFL/HAA
S-KO
HT-RBPJ WT
Propability
Dissociation rate (s⁻¹)

b)

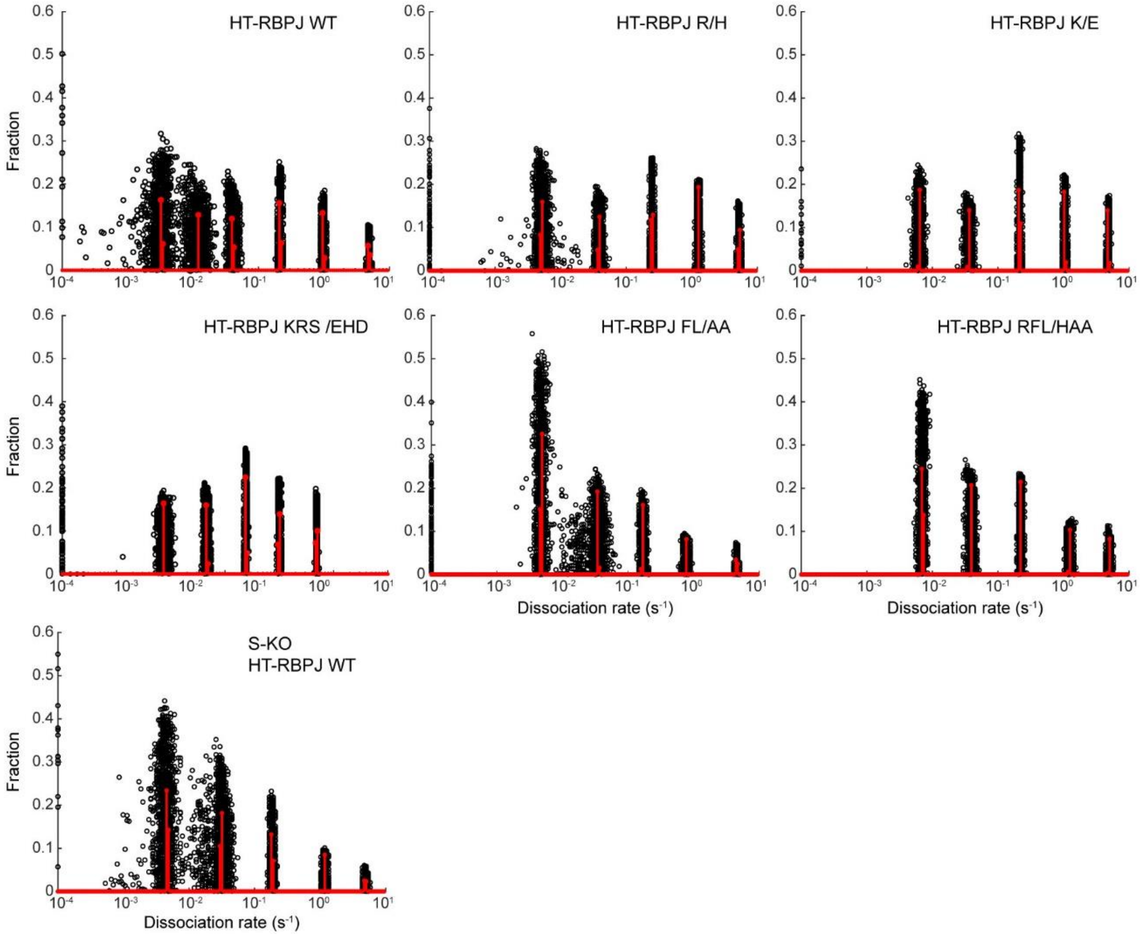
HT-RBPJ WT
HT-RBPJ R/H
HT-RBPJ K/E
HT-RBPJ KRS /EHD
HT-RBPJ FL/AA
HT-RBPJ RFL/HAA
S-KO
HT-RBPJ WT
Fraction
Dissociation rate (s⁻¹)

**Supplementary Figure 9: GRID dissociation rate spectra of HT-RBPJ variants**

**a)** Event and **b)** state spectrum of HT-RBPJ variants obtained with GRID. Resampling was performed by repeating GRID analysis for 499 times using 80% of data for each run. Experimental statistics are listed in Supplementary Table 3. Source data are provided as Source Data file for Supplementary Figure 9a,b.

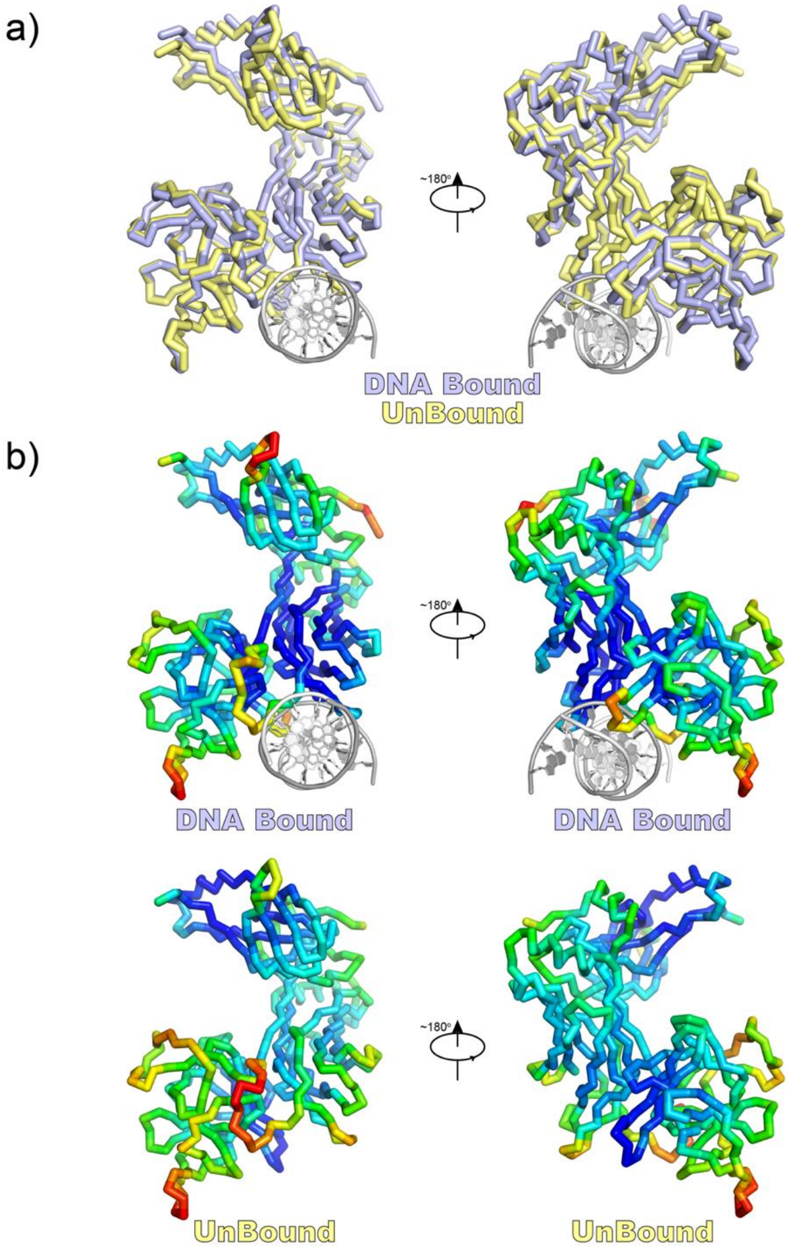

**Supplementary Figure 10 Structural Comparison of RBPJ/Su(H) in DNA-bound and -unbound states**

The figure shows a structural overlay and comparison of the two RBPJ/Su(H) molecules found in the asymmetric unit of the X-ray structure 5E24, in which one molecule is DNA-bound and the other is DNA-free [1].

**a)** Structural alignment of DNA-bound (light blue) and DNA-free (yellow) forms of RBPJ/Su(H), which highlights the conformational differences between the two states.

**b)** Temperature-factor comparison of the DNA-bound and DNA-free states of RBPJ/Su(H), in which higher and lower temperature factors are colored in a gradient from red to blue. When compared to the DNA-bound state, the DNA-free state of RBPJ/Su(H) has higher temperature-factors within its DNA binding domains.

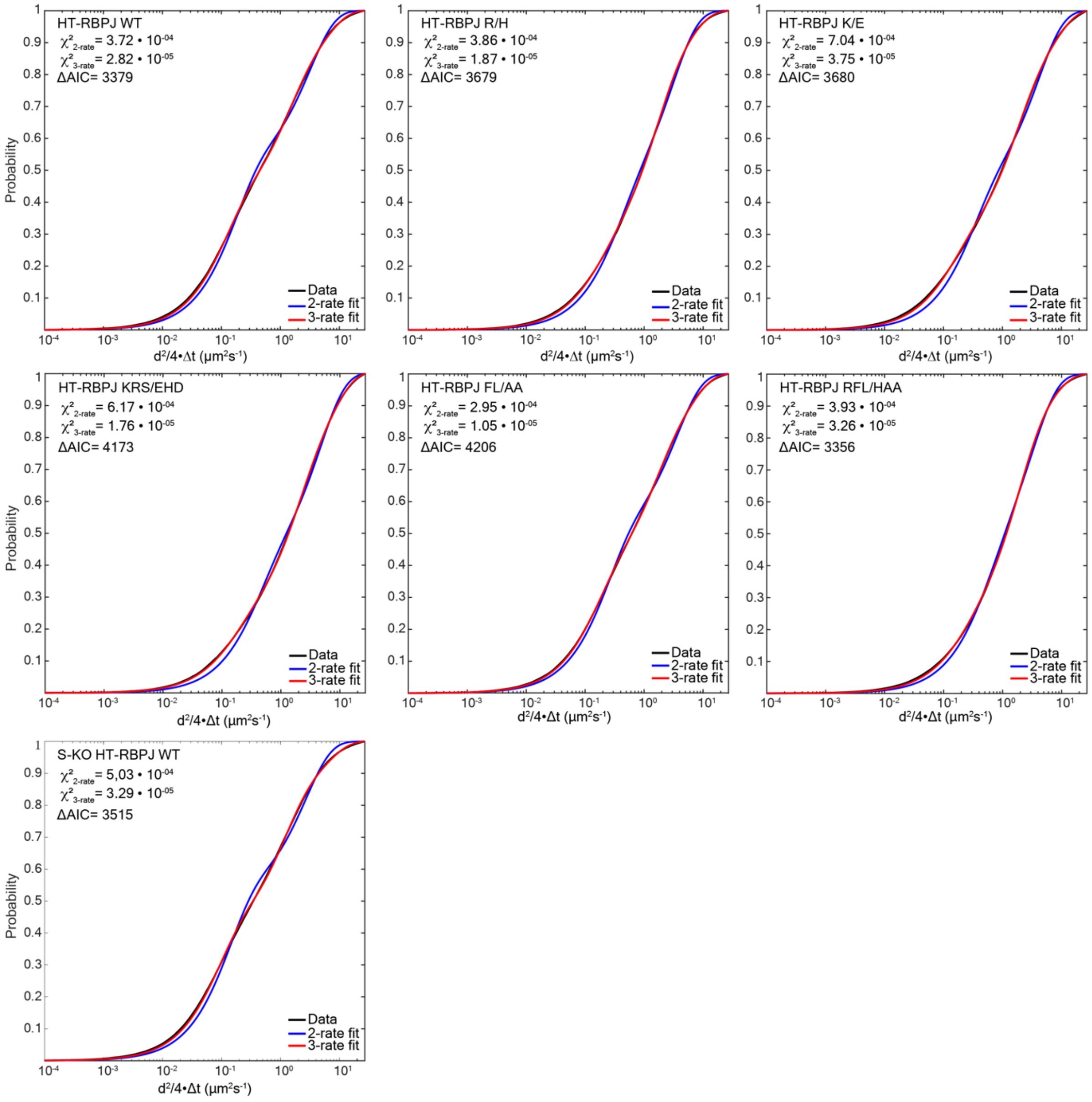

**Supplementary Figure 11: Analysis of diffusion measurements**

Cumulative jump distance distributions (black) of HT-RBPJ variants and fit of a two-component (blue) and a three-component diffusion model [2]. According to the reduced $\chi^2$ and the Akaike Information Criterion (AIC), the 3-component diffusion model described the data better. Experimental statistics are listed in Supplementary Table 5.

Source data are provided as Source Data file for Supplementary Figure 11.

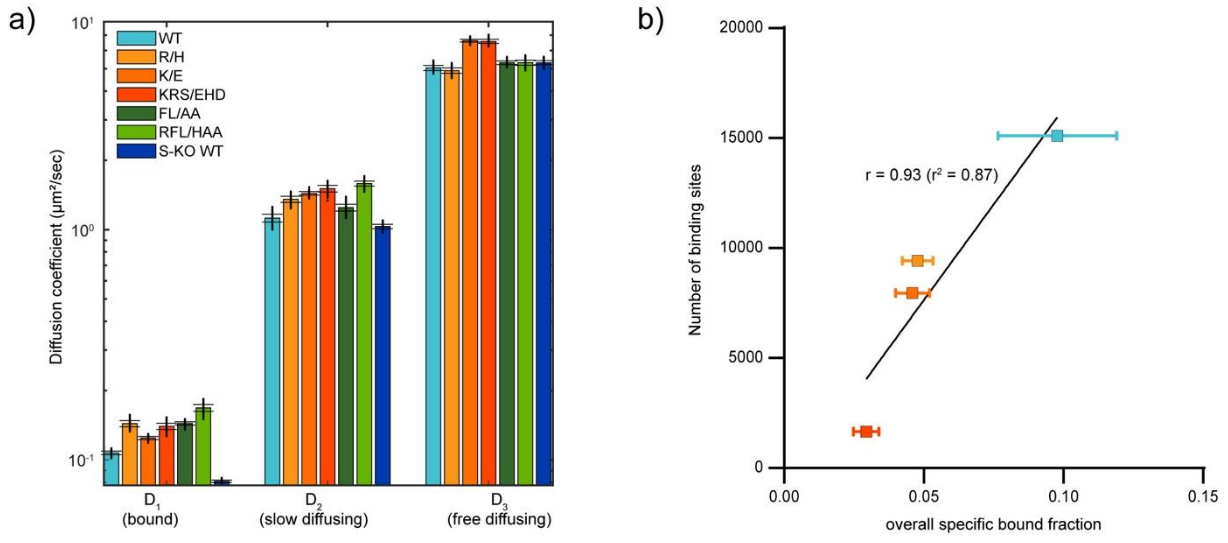

**Supplementary Figure 12: Diffusion coefficients and bound fractions of HT-RBPJ variants**

**a)** Diffusion coefficients of the three-component diffusion model and assignment to bound, slow and fast diffusing molecules. Data represents mean values ± s.d. from 400 resamplings with randomly selected 80 % of the data. Data are shown in Supplementary Table 4 and experimental statistics are listed in Supplementary Table 5.

**b)** Number of DNA binding sites found by ChIP-Seq versus overall specific bound fraction of HT-RBPJ variants. Pearson's correlation coefficient calculated for HT-RBPJ-WT and DNA binding mutants without considering variants with mutated cofactor binding interface (white squares).

Source data are provided as Source Data file for Supplementary Figure 12a,b.

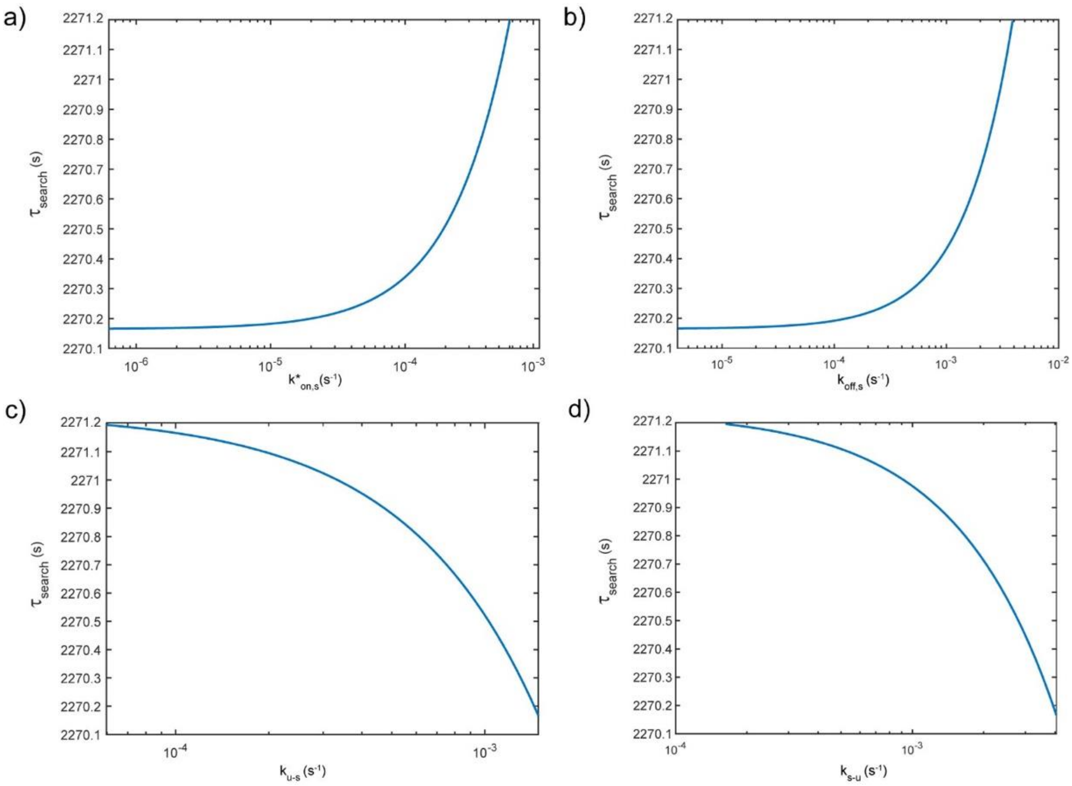

**Supplementary 13: Target site search time as function of various parameters of the three-state model of facilitated diffusion.**

Target site search time as function of **a)** direct association rate $k^*_{on,s}$, **b)** direct dissociation rate $k_{off,s}$, **c)** microscopic association rate $k_{u-s}$., and **d)** microscopic dissociation rate $k_{s-u}$. Data are provided as Source Data file for Supplementary Figure 13.

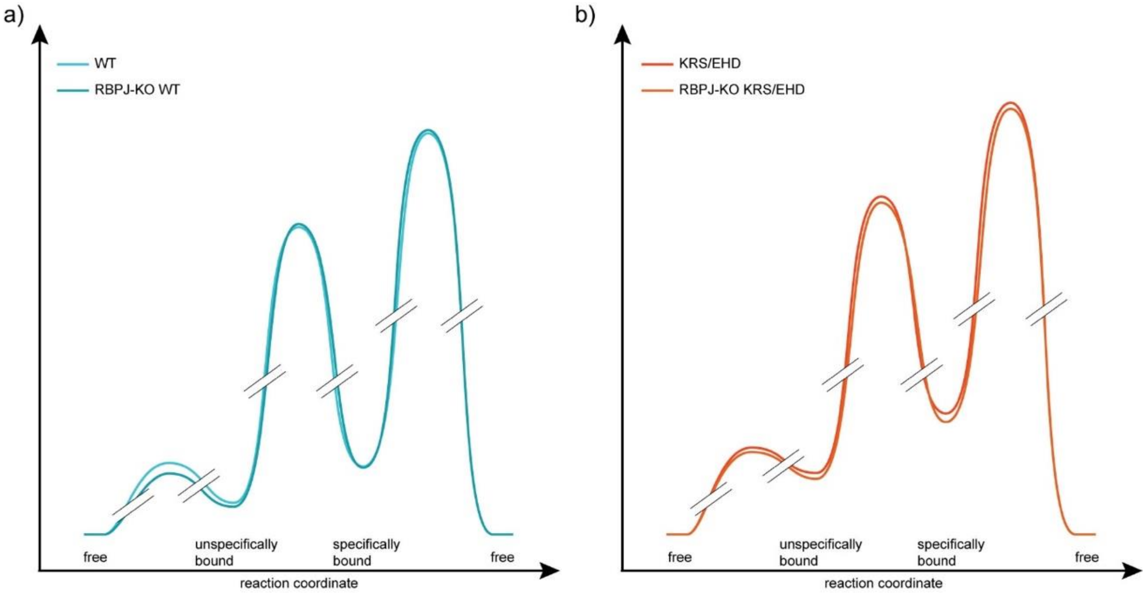

**Supplementary Figure 14: Comparison of HT-RBPJ binding energy landscapes in presence and absence of endogenous RBPJ**

Binding energy landscape of **a)** HT-RBPJ-WT and **b)** the triple DNA mutant HT-RBPJ-KRS/EHD in HeLa cells and in HeLa RBPJ knock out (#42) cells. Data are summarized in Supplementary Table 8. Data are provided as Source Data file for Supplementary Figure 14.

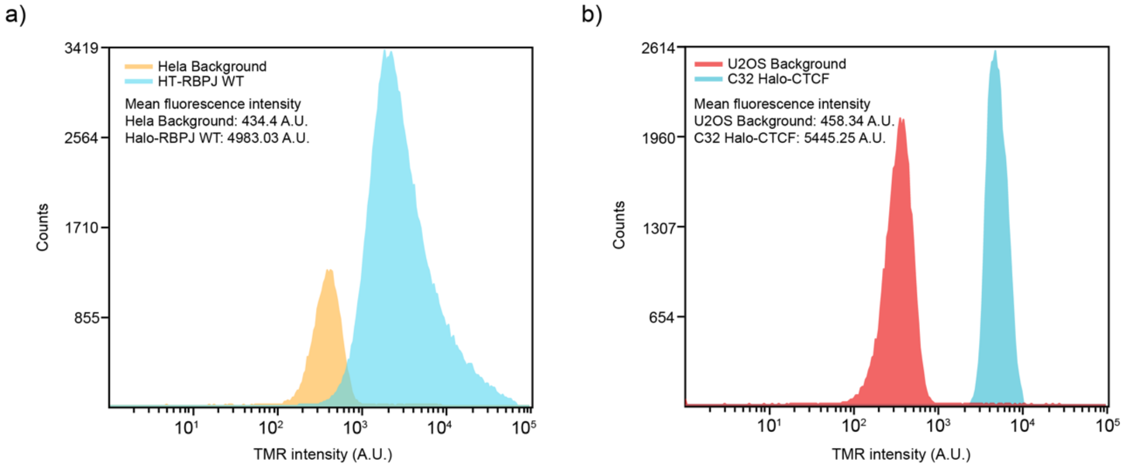

**Supplementary Figure 15: Quantification of HT-RBPJ-WT molecule abundance**

**a)** Flow cytometry intensity counts of HeLa cells stably expressing HT-RBPJ-WT labeled with HTL-TMR. **b)** Flow cytometry intensity counts of U2OS C32 HT-CTCF cells labeled with HTL-TMR were used as reference (Cattoglio et al. 2019). Unstained HeLa or U2OS cells were used to estimate the background. Number of Events for HeLa unstained/Halo-RBPJ-WT TMR-stained: 25762/133444. Number of Events for U2OS unstained/ Halo-CTCF TMR-stained: 5927/38979.

Supplementary Tables

| | | | | Fractions of event spectrum | | Fractions of state spectrum | |
|---|---|---|---|---|---|---|---|
| | **Variant** | **$k_d$ ($s^{-1}$)** | **$k_{off,u}$ ($s^{-1}$)** | **$A_s^e$ (%)** | **$A_u^e$ (%)** | **$A_s^s$ (%)** | **$A_u^s$ (%)** |
| **HeLa** | **HT-RBPJ-WT** | 0.00405 ± 0.0013 | 0.907 ± 0.04 | 0.17 ± 0.08 | 99.8 ± 0.1 | 26.7 ± 5.8 | 72.6 ± 6.7 |
| | **HT-RBPJ-R/H** | 0.0051 ± 0.001 | 1.42 ± 0.06 | 0.12 ± 0.02 | 99.9 ± 0.02 | 24.4 ± 4.3 | 70.4 ± 7.1 |
| | **HT-RBPJ-K/E** | 0.0063 ± 0.0006 | 1.28 ± 0.05 | 0.12 ± 0.02 | 99.9 ± 0.02 | 19.6 ± 2.6 | 80.0 ± 2.5 |
| | **HT-RBPJ-KRS/EHD** | 0.0076 ± 0.001 | 1.35 ± 0.03 | 0.11 ± 0.02 | 99.9 ± 0.02 | 16.3 ± 2.2 | 81.9 ± 5.0 |
| | **HT-RBPJ-FL/AA** | 0.0047 ± 0.0006 | 0.69 ± 0.04 | 0.60 ± 0.09 | 99.4 ± 0.1 | 45.6 ± 4.7 | 51.5 ± 4.4 |
| | **HT-RBPJ-RFL/HAA** | 0.007 ± 0.0006 | 1.01 ± 0.05 | 0.43 ± 0.06 | 99.6 ± 0.06 | 38.2 ± 2.9 | 61.8 ± 2.9 |
| **HeLa S-KO** | **HT-RBPJ-WT** | 0.0044 ± 0.001 | 0.6 ± 0.05 | 0.42 ± 0.11 | 99.6 ± 0.11 | 36.2 ± 5.3 | 62.7 ± 5.8 |
| **HeLa RBPJ-KO** | **HT-RBPJ-WT** | 0.0039 ± 0.0003 | 0.93 ± 0.03 | 0.20 ± 0.02 | 99.8 ± 0.019 | 31.0 ± 2.9 | 63.7 ± 3.8 |
| **HeLa RBPJ-KO** | **HT-RBPJ-KRS/EHD** | 0.00702 ± 0.0014 | 1.3 ± 0.054 | 0.115 ± 0.027 | 99.9 ± 0.027 | 16.9 ± 3.5 | 80.5 ± 5.8 |

**Supplementary Table 1**: Dissociation rates and amplitudes of state and event spectra obtained by GRID analysis of survival time distributions shown in Figure 2b and Supplementary Figure 8b.

| | **Variant** | **Specific residence time ($\tau_s$) ± s.d. (s)** | **Unspecific residence time ($\tau_u$) ± s.d. (s)** |
|---|---|---|---|
| **HeLa** | **Halo-RBPJ-WT** | 246.9 ± 79.3 | 1.1 ± 0.09 |
| | **Halo-RBPJ-R/H** | 194.2 ± 26.0 | 0.71 ± 0.02 |
| | **Halo-RBPJ-K/E** | 159.2 ± 14.7 | 0.78 ± 0.03 |
| | **Halo-RBPJ-KRS/EHD** | 133.5 ± 17.0 | 0.74 ± 0.02 |
| | **Halo-RBPJ-FL/AA** | 211.9 ± 25.9 | 1.43 ± 0.09 |
| | **Halo-RBPJ RFL/HAA** | 142.3 ± 11.7 | 0.99 ± 0.05 |
| **HeLa S-KO** | **Halo-RBPJ-WT** | 226.8 ± 50.9 | 1.67 ± 0.13 |
| **HeLa RBPJ-KO** | **HT-RBPJ-WT** | 277.8 ± 108.0 | 0.92 ± 0.08 |
| **HeLa RBPJ-KO** | **HT-RBPJ-KRS/EHD** | 142.45 ± 28.4 | 0.77 ± 0.03 |

**Supplementary Table 2**: Residence times of Halo-RBPJ variants at specific and unspecific sites, shown in Figure 2e and f.

| | | HeLa | | | | | | HeLa S-KO | HeLa RBPJ-KO | HeLa RBPJ-KO |
|---|---|---|---|---|---|---|---|---|---|---|
| **time-lapse condi-tion (s)** | | **HT-RBPJ-WT** | **HT-RBPJ-R/H** | **HT-RBPJ-K/E** | **HT-RBPJ-KRS/EHD** | **HT-RBPJ-FL/AA** | **HT-RBPJ-RFL/HAA** | **HT-RBPJ-WT** | **HT-RBPJ-WT** | **HT-RBPJ-KRS/EHD** |
| **0.1** | **#movies** | 21 | 19 | 19 | 26 | 27 | 34 | 11 | 27 | 20 |
| | **#tracks** | 1459 | 1329 | 1039 | 1904 | 669 | 621 | 723 | 2167 | 1283 |
| | **#all events** | 19835 | 16990 | 17073 | 25937 | 14015 | 19846 | 9599 | 28193 | 13788 |
| **0.4** | **#movies** | 15 | 27 | 21 | 26 | 30 | 29 | 24 | 27 | 18 |
| | **#tracks** | 889 | 1064 | 1135 | 1669 | 731 | 402 | 981 | 2089 | 1013 |
| | **#all events** | 15220 | 20562 | 23255 | 26965 | 15747 | 17694 | 17988 | 34393 | 15373 |
| **1.6** | **#movies** | 28 | 24 | 14 | 26 | 23 | 37 | 14 | 31 | 17 |
| | **#tracks** | 2648 | 1978 | 1194 | 2711 | 1148 | 1024 | 974 | 4540 | 1431 |
| | **#all events** | 26782 | 22143 | 14678 | 31853 | 16386 | 20004 | 11724 | 48701 | 14117 |
| **6.4** | **#movies** | 27 | 20 | 20 | 18 | 27 | 24 | 19 | 18 | 24 |
| | **#tracks** | 1446 | 882 | 920 | 856 | 966 | 548 | 890 | 1603 | 864 |
| | **#all events** | 17315 | 11619 | 13110 | 12744 | 11931 | 9436 | 9617 | 1603 | 12410 |

**Supplementary Table 3:** Statistics of single-molecule time-lapse measurements.

| | **Variant** | **$D_1$ (µm²/s)** | **$D_2$ (µm²/s)** | **$D_3$ (µm²/s)** | **$A_1$ (bound)** | **$A_2$ (slow diffusing)** | **$A_3$ (free diffusing)** |
|---|---|---|---|---|---|---|---|
| **HeLa** | **Halo-RBPJ-WT** | 0.107 ± 0.001 | 1.12 ± 0.02 | 5.0 ± 0.1 | 0.37 ± 0.01 | 0.35 ± 0.01 | 0.29 ± 0.01 |
| | **Halo-RBPJ-R/H** | 0.144 ± 0.002 | 1.36 ± 0.02 | 4.9 ± 0.1 | 0.20 ± 0.01 | 0.49 ± 0.01 | 0.31 ± 0.01 |
| | **Halo-RBPJ-K/E** | 0.125 ± 0.001 | 1.44 ± 0.01 | 6.6 ± 0.1 | 0.23 ± 0.01 | 0.44 ± 0.01 | 0.33 ± 0.01 |
| | **Halo-RBPJ-KRS/EHD** | 0.140 ± 0.002 | 1.51 ± 0.01 | 6.6 ± 0.1 | 0.18 ± 0.01 | 0.40 ± 0.01 | 0.41 ± 0.01 |
| | **Halo-RBPJ-FL/AA** | 0.144 ± 0.001 | 1.24 ± 0.02 | 5.3 ± 0.1 | 0.33 ± 0.01 | 0.35 ± 0.01 | 0.32 ± 0.01 |
| | **Halo-RBPJ-RFL/HAA** | 0.169 ± 0.003 | 1.59 ± 0.02 | 5.3 ± 0.1 | 0.15 ± 0.01 | 0.56 ± 0.01 | 0.29 ± 0.01 |
| **HeLa S-KO** | **Halo-RBPJ-WT** | 0.081 ± 0.001 | 1.03 ± 0.01 | 5.3 ± 0.1 | 0.38 ± 0.01 | 0.40 ± 0.01 | 0.22 ± 0.01 |
| **Hela RBPJ-KO** | **HT-RBPJ-WT** | 0.124 ± 0.001 | 1.21 ± 0.01 | 6.2 ± 0.03 | 0.39 ± 0.01 | 0.40 ± 0.01 | 0.22 ± 0.01 |
| **Hela RBPJ-KO** | **HT-RBPJ-KRS/EHD** | 0.1 ± 0.01 | 1.22 ± 0.01 | 6.2 ± 0.03 | 0.21 ± 0.01 | 0.43 ± 0.02 | 0.36 ± 0.01 |

**Supplementary Table 4:** Diffusion coefficients and fractions obtained from cumulative jump distance distributions shown in Figure 3c and Supplementary Figure 9.

| | Variant | Days | #Cells | #Tracks | #all events |
|---|---|---|---|---|---|
| **HeLa** | **Halo-RBPJ-WT** | 4 | 9 | 25204 | 57546 |
| | **Halo-RBPJ-R/H** | 2 | 12 | 26237 | 55238 |
| | **Halo-RBPJ-K/E** | 2 | 7 | 77936 | 174652 |
| | **Halo-RBPJ-KRS/EHD** | 2 | 11 | 27630 | 60785 |
| | **Halo-RBPJ-FL/AA** | 3 | 19 | 33548 | 77632 |
| | **Halo-RBPJ-RFL/HAA** | 3 | 22 | 55242 | 110146 |
| **HeLa S-KO** | **Halo-RBPJ-WT** | 1 | 13 | 42339 | 100670 |
| **Hela RBPJ-KO** | **HT-RBPJ WT** | 2 | 25 | 167314 | 366421 |
| **Hela RBPJ-KO** | **HT-RBPJ-KRS** | 2 | 24 | 147086 | 315285 |

**Supplementary Table 5**: Statistics of continuous movies for diffusion analysis.

| | $k_{on,u}$ ($s^{-1}$) | $k_{u-s}$ ($s^{-1}$) | $k_{s-u}$ ($s^{-1}$) | $k_{off,s}$ ($s^{-1}$) | $k_{on,s}$ ($s^{-1}$) | $\tau_{search}$ (s) |
|---|---|---|---|---|---|---|
| **Max_$\tau_{search}$** | 0.3806 ± 0.0813 | 0.0004 ± 0.0001 | 0.0012 ± 0.0006 | 0.0029 | 0.0004 ± 0.0002 | 2270.9 ± 616.5 |
| **Min_$\tau_{search}$** | 0.3806 ± 0.0813 | 0.0015 ± 0.0003 | 0.0041 ± 0.0019 | $4.05*10^{-6}$ | $6.25*10^{-7}$ ± $2.73*10^{-7}$ | 2270.1 ± 582.0 |

**Supplementary Table 6:** Limits of the parameters of the three-state model of facilitated diffusion determined for HT-RBPJ-WT. Errors were calculated based on Gaussian error propagation. $k_{off,s}$ was fixed relative to $k_d$.

| | Variant | $p_s$ | $p_u$ | $p_f$ | $k_{on,u}$ ($s^{-1}$) | $k_{off,u}$ ($s^{-1}$) | $k_{u-s}$ ($s^{-1}$) | $k_{s-u}$ ($s^{-1}$) | $k_d$ ($s^{-1}$) | $k_{off,s}$ ($s^{-1}$) | $k_{on,s}$ ($s^{-1}$) | $\tau_{search}$ (s) |
|---|---|---|---|---|---|---|---|---|---|---|---|---|
| **HeLa** | **HT-RBPJ-WT** | 0.10 ± 0.02 | 0.27 ± 0.03 | 0.63 ± 0.01 | 0.3806 ± 0.0813 | 0.907 ± 0.07 | 0.0013 ± 0.0003 | 0.0038 ± 0.0018 | 0.0041 ± 0.0013 | 0.000286 | 0.0000441 ± 0.0000166 | 2270.2 ± 538.3 |
| | **HT-RBPJ-R/H** | 0.05 ± 0.01 | 0.14 ± 0.01 | 0.80 ± 0.01 | 0.2639 ± 0.0696 | 1.41 ± 0.04 | 0.0015 ±0.0003 | 0.0048 ± 0.0027 | 0.0052 ± 0.001 | 0.000363 | 0.0000217 ± 0.0000100 | 3854.3 ± 1073.5 |
| | **HT-RBPJ-K/E** | 0.05 ± 0.01 | 0.19 ± 0.01 | 0.77 ± 0.01 | 0.3139 ± 0.0493 | 1.28 ± 0.05 | 0.0014 ± 0.0003 | 0.0058 ± 0.0029 | 0.0063 ± 0.0006 | 0.000443 | 0.0000266 ± 0.0000115 | 3299.8 ± 733.3 |
| | **HT-RBPJ-KRS/EHD** | 0.03 ± 0.01 | 0.15 ± 0.01 | 0.82 ± 0.01 | 0.2491 ± 0.0803 | 1.35 ± 0.04 | 0.0014 ± 0.0003 | 0.007 ± 0.0055 | 0.0075 ± 0.001 | 0.000528 | 0.0000190 ± 0.0000133 | 4391.5 ± 1385.7 |
| | **HT-RBPJ-FL/AA** | 0.15 ± 0.02 | 0.17 ± 0.01 | 0.67 ± 0.01 | 0.1786 ± 0.0373 | 0.70 ± 0.04 | 0.0039 ± 0.0008 | 0.0044 ± 0.0016 | 0.0047 ± 0.0006 | 0.000333 | 0.0000763 ± 0.0000147 | 1159.6 ± 282.7 |
| | **HT-RBPJ-RFL/HAA** | 0.06 ± 0.01 | 0.09 ± 0.01 | 0.85 ± 0.01 | 0.1120 ± 0.0446 | 1.01 ± 0.05 | 0.0041 ± 0.0008 | 0.0066 ± 0.0041 | 0.007 ± 0.0006 | 0.000496 | 0.0000340 ± 0.0000148 | 2305.0 ± 881.2 |
| **HeLa S-KO** | **HT-RBPJ-WT** | 0.14 ± 0.02 | 0.24 ± 0.02 | 0.62 ± 0.01 | 0.2289 ± 0.0444 | 0.6 ± 0.05 | 0.0024 ± 0.0005 | 0.0041 ± 0.0015 | 0.0044 ± 0.001 | 0.000311 | 0.0000691 ± 0.0000156 | 1408.5 ± 324.3 |
| **Hela RBPJ -KO** | **HT-RBPJ-WT** | 0.10 ± 0.02 | 0.29 ± 0.03 | 0.61 ± 0.01 | 0.5086 ± 0.1098 | 1.09 ± 0.10 | 0.0012 ± 0.0002 | 0.0034 ± 0.0017 | 0.0036 ± 0.0014 | 0.000254 | 0.0000405 ± 0.0000160 | 2552.9 ± 602.3 |
| **Hela RBPJ -KO** | **HT-RBPJ-KRS/EHD** | 0.04 ± 0.01 | 0.17 ± 0.01 | 0.79 ± 0.01 | 0.2815 ± 0.0857 | 1.3 ± 0.05 | 0.0014 ± 0.0003 | 0.0065 ± 0.0056 | 0.0070 ± 0.0014 | 0.000495 | 0.0000225 ± 0.0000173 | 3810.8 ± 1147.0 |

**Supplementary Table 7:** Overall fractions, kinetic rates and target site search time for RBPJ variants shown in Figure 3e. Errors were calculated by Gaussian error propagation. Error for $k_{u-s}$ was estimated with 20% of the value. $k_{off,s}$ was fixed relative to $k_d$.

| | Variant | $\Delta G^{\#}_{f\text{->}u}$ ($k_BT$) | $\Delta G_u$ ($k_BT$) | $\Delta G^{\#}_{u\text{->}s}$ ($k_BT$) | $\Delta G_s$ ($k_BT$) | $\Delta G^{\#}_{f\text{->}s}$ ($k_BT$) |
|---|---|---|---|---|---|---|
| **HeLa** | **Halo-RBPJ-WT** | 0.97 ± 0.21 | 0.87 ± 0.23 | 7.45 ± 1.49 | 1.87 ± 0.38 | 10.03 ± 0.38 |
| | **Halo-RBPJ-R/H** | 1.33 ± 0.26 | 1.68 ± 0.27 | 8.16 ± 1.63 | 2.82 ± 0.46 | 10.74 ± 0.46 |
| | **Halo-RBPJ-K/E** | 1.16 ± 0.16 | 1.41± 0.16 | 7.96 ± 1.59 | 2.81 ± 0.43 | 10.53 ± 0.43 |
| | **Halo-RBPJ-KRS/EHD** | 1.39 ± 0.32 | 1.69 ± 0.32 | 8.29 ± 1.66 | 3.32 ± 0.70 | 10.87 ± 0.70 |
| | **Halo-RBPJ-FL/AA** | 1.72 ± 0.21 | 1.36 ± 0.22 | 6.90 ± 1.38 | 1.47 ± 0.19 | 9.48 ± 0.19 |
| | **Halo-RBPJ-RFL/HAA** | 2.19 ± 0.40 | 2.20 ± 0.40 | 7.71 ±1.54 | 2.68 ± 0.44 | 10.29 ± 0.44 |
| **HeLa S-KO** | **Halo-RBPJ-WT** | 1.48 ± 0.20 | 0.96 ± 0.21 | 7.00 ±1.40 | 1.50 ± 0.23 | 9.58 ± 0.24 |
| **Hela RBPJ-KO** | **HT-RBPJ-WT** | 0.68 ± 0.22 | 0.76 ± 0.23 | 7.54 ± 1.51 | 1.84 ± 0.40 | 10.12 ± 0.40 |
| **Hela RBPJ-KO** | **HT-RBPJ-KRS/EHD** | 1.27 ± 0.31 | 1.53 ± 0.31 | 8.12 ± 1.62 | 3.09 ± 0.77 | 10.70 ± 0.77 |

**Supplementary Table 8:** Free energy differences between the free state and transition or bound states of HT-RBPJ variants shown in Figure 4e. Errors were calculated based on Gaussian error propagation.

| | Variant | $\Delta G^{\#}_{f\text{->}u}$ ($k_BT$) | $\Delta G_u$ ($k_BT$) | $\Delta G^{\#}_{u\text{->}s}$ ($k_BT$) | $\Delta G_s$ ($k_BT$) | $\Delta G^{\#}_{f\text{->}s}$ ($k_BT$) |
|---|---|---|---|---|---|---|
| **U2OS** | **TALE 9R** | 0.55 ± 0.40 | 0.42 ± 0.41 | 7.45 ± 1.49 | 1.90 ± 0.45 | 9.97 ± 0.45 |
| | **TALE 13R** | 0.97 ± 0.32 | 0.78 ± 0.32 | 6.46 ± 1.29 | 0.82 ± 0.34 | 8.98 ± 0.34 |
| | **TALE 15R** | 1.07 ± 0.33 | 0.69 ± 0.34 | 7.05 ± 1.41 | 0.96 ± 0.30 | 9.56 ± 0.30 |
| | **TALE 19R** | 0.41 ± 0.27 | 0.61 ± 0.28 | 6.80 ± 1.36 | 1.50 ± 0.40 | 9.32 ± 0.40 |

**Supplementary Table 9:** Free energy differences between the free state and transition or bound states of TALE-TFs shown in Figure 4f. Errors were calculated based on Gaussian error propagation.

| | Variant | $\Delta G^{\#}_{f\text{->}u}$ ($k_BT$) | $\Delta G_u$ ($k_BT$) | $\Delta G^{\#}_{f\text{->}s}$ ($k_BT$) | $\Delta G_s$ ($k_BT$) |
|---|---|---|---|---|---|
| **Embryonic stem cells** | **SOX2** | 1.30 ± 0.21 | 1.51 ± 0.15 | 5.93 ± 0.05 | 3.44 ± 0.16 |
| | **SOX2D** | 0.84 ± 0.17 | 1.04 ± 0.21 | 5.14 ± 0.09 | 2.98 ± 0.15 |
| | **SOX2M** | 1.72 ± 0.17 | 1.97 ± 0.22 | 6.40 ± 0.09 | 4.17 ± 0.23 |
| **Hemocytes** | **GAF** | 0.95 | -0.36 | 5.00 | 0.14 |
| | **GAF dPOZ** | 0.28 | -0.62 | 4.65 | 0.89 |
| | **GAF dQ** | 0.54 | -0.53 | 4.35 | 0.46 |

**Supplementary Table 10:** Free energy differences between the free state and transition or bound states shown in figure 4g and 4h calculated from published kinetic rates. Errors were calculated based on Gaussian error propagation.

| Plasmids | Description |
|---|---|
| pGa981-6 | Reporter plasmid containing twelve RBPJ binding sites, a Beta-Globin promoter as well as the gene for Luciferase |
| pcDNA3.1(-)hsNICD | Expression plasmid for human Notch1 Intracellular domain |
| pcDNA3.1-Flag-mRBPJ-CRr(wt) | Expression plasmid for murine Flag-tagged RBPJ wildtype |
| pcDNA3.1-Flag-mRBPJ-CRr(R218H) | Expression plasmid for murine Flag-tagged RBPJ R218H |
| pcDNA3.1-Flag-mRBPJ-CRr(K195E) | Expression plasmid for murine Flag-tagged RBPJ K195E |
| pcDNA3.1-Flag-mRBPJ-CRr(KRS) | Expression plasmid for murine Flag-tagged RBPJ K195E/R218H/S221D |
| pcDNA3.1-Flag-mRBPJ-CRr(F261A/L388A) | Expression plasmid for murine Flag-tagged RBPJ F261A/L388A |
| pcDNA3.1-Flag-mRBPJ-CRr(R218H/F261A/L388A) | Expression plasmid for murine Flag-tagged RBPJ R218H/F261A/L388A |
| pcDNA3.1-Flag-mRBPJ-CRr-VP16(wt) | Expression plasmid for murine Flag-tagged and VP16-fused RBPJ wildtype |
| pcDNA3.1-Flag-mRBPJ-CRr-VP16(R218H) | Expression plasmid for murine Flag-tagged and VP16-fused RBPJ R218H |
| pcDNA3.1-Flag-mRBPJ-CRr-VP16(K195E) | Expression plasmid for murine Flag-tagged and VP16-fused RBPJ K195E |
| pcDNA3.1-Flag-mRBPJ-CRr-VP16(KRS) | Expression plasmid for murine Flag-tagged and VP16-fused RBPJ KRS |
| pcDNA3.1-Flag-mRBPJ-CRr-VP16(F261A/L388A) | Expression plasmid for murine Flag-tagged and VP16-fused RBPJ F261A/L388A |
| pcDNA3.1-Flag-mRBPJ-CRr-VP16(R218H/F261A/L388A) | Expression plasmid for murine Flag-tagged and VP16-fused RBPJ R218H/F261A/L388A |

**Supplementary Table 11:** List of plasmids used in luciferase assays shown in Figure 1b and 1c.

| Plasmids | Description |
|---|---|
| pLVTO-Halo-mRBPJ-CRr(wt) | Expression plasmid for murine Halo-tagged RBPJ wildtype |
| pLVTO-Halo-mRBPJ-CRr(KRS) | Expression plasmid for murine Halo-tagged RBPJ K195E/R218H/S221D |
| pcDNA3.1-EGFP-mRBPJ-CRr(wt) | Expression plasmid for murine EGFP-tagged RBPJ wildtype |
| pcDNA3.1-EGFP-mRBPJ-CRr(KRS) | Expression plasmid for murine EGFP-tagged RBPJ K195E/R218H/S221D |
| pcDNA3.1(-)Flag1-hsNICD | Expression plasmid for human Flag-tagged Notch1 Intracellular domain |

**Supplementary Table 12:** List of plasmids used in immunoprecipitation shown in Figure S2a

| CRISPR/Cas9 guides | |
|---|---|
| hRBPJ guide #1 | 5’- TCA TGC CAG TTC ACA GCA GT -3’ |
| hRBPJ guide #2 | 5’- TCC TTC TAC ATG CAA GTA TC -3’ |
| hSHARP guide #1 | 5’- GCA ACT TAC CCG AGA ACG TG -3’ |
| hSHARP guide #2 | 5´- TCC CGG TGA ATC AAA GAC GG -3’ |

**Supplementary Table 13:** Guide sequences used for CRISPR/Cas9 experiments to generate knock-out cell lines.

| Gene | Primer | Gene Globe ID |
|---|---|---|
| *ACTIN* | Hs_ACTB_2_SG QuantiTect Primer Assay | QT01680476 |
| *HES1* | Hs_HES1_1_SG QuantiTect Primer Assay | QT00039648 |
| *HEY1* | Hs_HEY1_1_SG QuantiTect Primer Assay | QT00035644 |
| *HEY2* | Hs_HEY2_3_SG QuantiTect Primer Assay | QT01849701 |
| *SHARP* | Hs_SPEN_1_SG QuantiTect Primer Assay | QT00042196 |

**Supplementary Table 14:** Primers used for real time qPCR.

| Antibody | Catalogue Nr.: | Company | Type |
|---|---|---|---|
| anti-RBPJ | 5313 | Cell Signaling Technol-ogy | monoclonal (rabbit) |

**Supplementary Table 15:** Antibody used in ChIP-seq experiments

| Primary Antibody | Catalogue Nr.: | Company | Type | Suppl. Figure |
|---|---|---|---|---|
| Anti-RBPJ (T6709) | IIM-2ARBP2 | Cosmo Bio Co., Ltd. | Monoclonal (Rat) | S2a, S3b, S4g, S7a, S7c |
| Anti-Beta-ACTIN (A1978) | A1978 | Sigma-Aldrich | Monoclonal (Mouse) | S3b, S4g, S4h, S7a, S7c |
| Anti-Flag (M5) | F4042 | Sigma-Aldrich | Monoclonal (Mouse) | S2a |
| Anti-SHARP.1 | - | - | Polyclonal antise-rum (Rabbit) | S4b |
| Anti-Halo-Tag (G921A) | G9211 | Promega GmbH | Monoclonal (Mouse) | S4h |
| **Secondary Antibody** | **Catalogue Nr.:** | **Company** | **Information** | **Suppl. Figure** |
| Peroxidase AffiniPure Anti-rat IgG HRP-conjugated | 112-035-071 | Jackson Immu-noResearch | Whole antibody | S3b, S4g, S7a, S7c |
| ECL™ Anti-mouse IgG HRP-conju-gated | NA931V | Cytiva | Whole antibody | S3b, S4g, S4h, S7a, S7c |
| Rabbit IgG HRP Linked Whole Ab | NA934V | Cytiva | Whole antibody | S4b |

**Supplementary Table 16:** List of antibodies used for Western Blots.

| Primary Antibody | Catalogue Nr.: | Company | Type | Suppl. Figure |
|---|---|---|---|---|
| Anti-SHARP.1 | - | - | Polyclonal rabbit an-tiserum | S4c |
| **Secondary Antibody** | **Catalogue Nr.:** | **Company** | **Type** | **Suppl. Figure** |
| Goat anti-Rabbit IgG (H+L) Cross-Adsorbed secondary Antibody, Alexa Fluor™ 488 | A11008 | Invitrogen | Polyclonal | S4c |

**Supplementary Table 17:** List of antibodies used for immunofluorescence microscopy.

### Supplementary Movie Legends

**Filename: Supplementary Movie 1**
Description: Example movie of HT-RBPJ-WT time-lapse imaging in a HeLa cell nucleus (dashed line) with 50 ms camera exposure time and 0.1 s frame cycle time. Related to Figure 2b. Scale bar: 4 µm.

**Filename: Supplementary Movie 2**
Description: Example movie of HT-RBPJ-WT imaging in a HeLa cell nucleus (dashed line) with continuous illumination at 11.7 ms frame cycle time. Scale bar: 4 µm.
